# Two H3K23 histone methyltransferases, SET-32 and SET-21, function synergistically to promote nuclear RNAi-mediated transgenerational epigenetic inheritance in *Caenorhabditis elegans*

**DOI:** 10.1101/2024.11.05.622152

**Authors:** Anna Zhebrun, Julie Z. Ni, Laura Corveleyn, Siddharth Ghosh Roy, Simone Sidoli, Sam G. Gu

## Abstract

Nuclear RNAi in *C. elegans* induces a set of transgenerationally heritable marks of H3K9me3, H3K23me3, and H3K27me3 at the target genes. The function of H3K23me3 in the nuclear RNAi pathway is largely unknown due to the limited knowledge of H3K23 histone methyltransferase (HMT). In this study we identified SET-21 as a novel H3K23 HMT. By taking combined genetic, biochemical, imaging, and genomic approaches, we found that SET-21 functions synergistically with a previously reported H3K23 HMT SET-32 to deposit H3K23me3 at the native targets of germline nuclear RNAi. We identified a subset of native nuclear RNAi targets that are transcriptionally activated in the *set-21;set-32* double mutant. SET-21 and SET-32 are also required for robust transgenerational gene silencing induced by exogenous dsRNA. The *set-21;set-32* double mutant strain exhibits an enhanced temperature-sensitive mortal germline phenotype compared to the *set-32* single mutant, while the *set-21* single mutant animals are fertile. We also found that HRDE-1 and SET-32 are required for cosuppression, a transgene-induced gene silencing phenomenon, in *C. elegans* germline. Together, these results support a model in which H3K23 HMTs SET-21 and SET-32 function cooperatively to ensure the robustness of germline nuclear RNAi and promotes the germline immortality under the heat stress.

## Introduction

RNA interference refers to a diverse set of gene silencing activities that are guided by the small interfering RNAs (siRNAs) ^1–3^. Broadly speaking, the underlying gene silencing mechanisms of RNAi fall into two categories: transcriptional gene silencing (TGS) ^4–8^ and post-transcriptional gene silencing (PTGS) ^9, 10^. The TGS mechanism, which is also referred to as nuclear RNAi, guides the formation of heterochromatin at transposons and other repetitive DNA, and plays an essential role in genome stability and germ cell development in plants, fungi, and animals ^11, 12^. Since its initial discovery in plants ^13, 14^ and *S. pombe* ^15^, nuclear RNAi has been used as a model system to explore different aspects of chromatin biology, particularly in the regulatory function of non-coding RNA and the mechanisms of transgenerational epigenetic inheritance (TEI) ^16, 17^.

In *C. elegans*, exogenous dsRNA or piRNA can induce various heterochromatin marks including H3K9me3 ^18, 19^, H3K27me3 ^20^, and H3K23me3 ^21^ at a target gene. Remarkably, RNAi-induced histone modifications and the silencing effect can persist for multiple generations ^18, 20, 21^, which makes *C. elegans* a tractable system to study TEI. The heterochromatic histone modifications also mark the native targets of the germline nuclear RNAi, which are largely composed of transposable elements ^20–23^. Surprisingly, although H3K9me3 is one of the best-known constitutive heterochromatin marks, we and others found that the H3K9me3 appears to be dispensable for transcriptional repression at the nuclear RNAi target genes ^24–26^. H3K27me3 is a hall mark for the facultative heterochromatin. In *C. elegans*, H3K27me2/3 in adult germ cells is deposited by the Drosphila E(Z) and human EZH2 homolog MES-2 ^27, 28^. Loss of MES-2 leads to sterility^28^, which makes it difficult to investigate the function of H3K27me3 in *C. elegans* nuclear RNAi.

Although H3K23me is an evolutionarily conserved heterochromatin mark found in plants ^29^, fungi ^30^, and animals including mammals ^30–37^, much less is known about this histone modification compared to H3K9me or H3K27me. Loss of H3K23me in *Tetrahymena* is associated with an increase in DNA damage ^32^ . H3K23me is an abundant histone modification throughout *C. elegans* development and is present in both the soma and germline^33, 34^. The whole-genome distributions of H3K23m3 and H3K9me3 are similar to each other, and both are highly enriched in the heterochromatin in *C. elegans* ^21^.

We previously reported that SET-32 can catalyze H3K23 methylation *in vitro* and is required for the nuclear RNAi-mediated H3K23me3 *in vivo* ^21^. However, Loss of SET-32 only leads to a partial loss of H3K23me3, indicating the existence of additional H3K23 HMTs^21^. Most of the known HTMs contain an evolutionarily conserved catalytic SET domain ^38^. In *C. elegans*, there are 38 SET domain-containing proteins ^39^. In this paper, we identified and characterized SET-21 as the other H3K23 HMT that functions in the germline nuclear RNAi pathway.

## Results

### SET-21 exhibits H3K23 methyltransferase activity *in vitro*

To identify other putative H3K23 methyltransferases, we performed a phylogenetic analysis of the 38 *C. elegans* SET domain-containing proteins^39^. Among them, SET-21 and SET-33 have the highest homology to SET-32 (Fig. 1A). *set-33* is listed as a pseudogene in the WormBase ^40^ and therefore was not investigated by this study.

**Figure 1.**
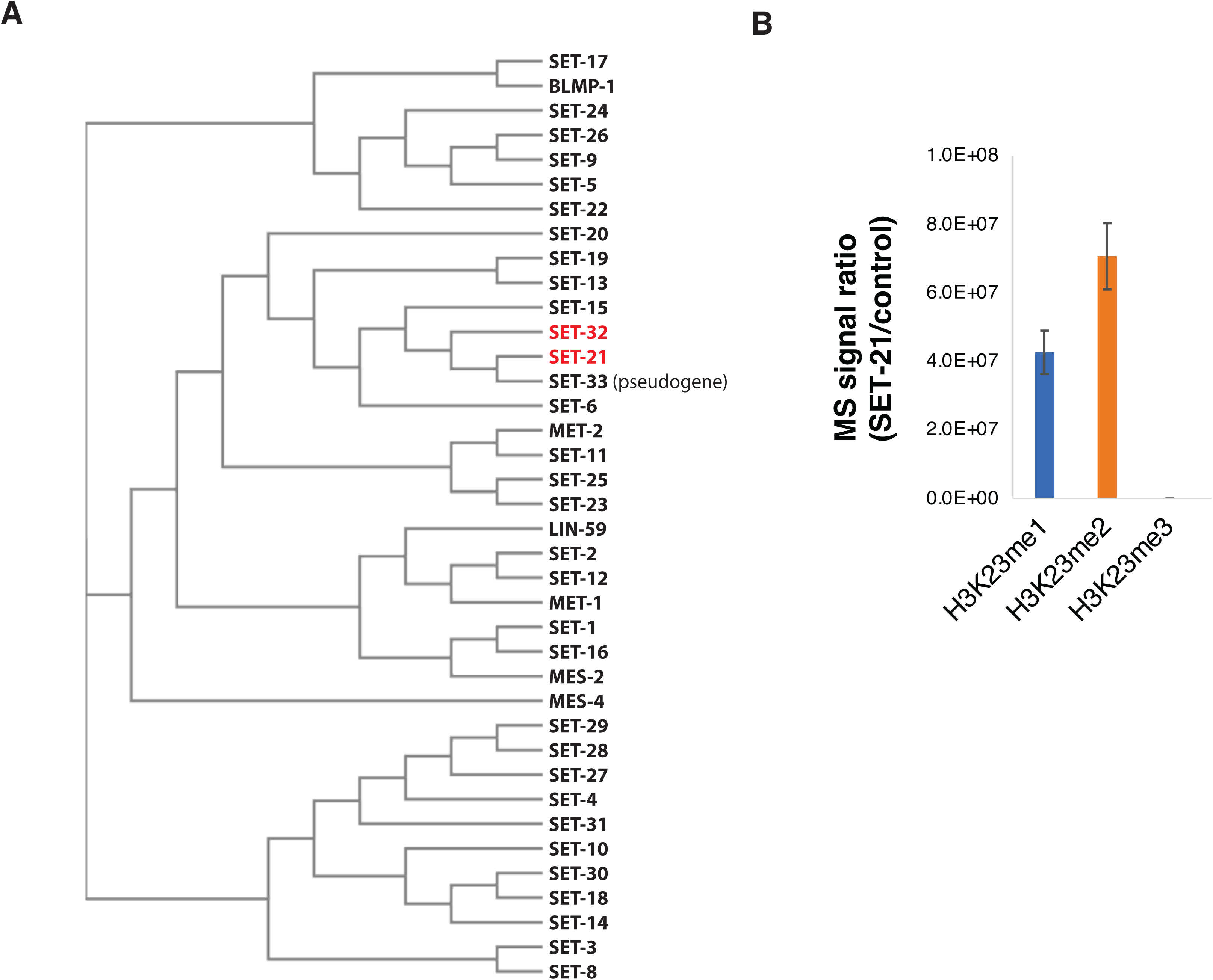
SET-21 is a H3K23 histone methyltransferase. (A) Phylogenetic tree of SET-domain-containing proteins in *C. elegans* ^39^ showing that SET-21 is the closest homolog of SET-32. (B) SET-21 methylates H3K23 *in vitro*. Mass spectrometry analysis of *in vitro* histone methyltransferase assay was performed by using recombinant GST-SET-21 or GST-3xFLAG (control) and recombinant *C. elegans* H3 proteins. The relative abundance of K23me1/2/3 for histone H3 peptide KQLATKAAR (aa 18–26) produced by GST-SET-21 or GST-3xFLAG were calculated and the ratios of the two (GST-SET-21/GST-3xFLAG) were plotted. Error bar: SEM. N=2 biological repeats.

SET-21 and SET-32 share 43% sequence identity, and the two genes also share similar gene structures (Fig. S1A, B). To determine whether SET-21 is an H3K23 HMT, we performed an *in vitro* HMT assay by using purified recombinant GST-SET-21 and H3 proteins. Mass spectrometry analysis of the reaction product showed that GST-SET-21 methylated the lysine 23 in H3. Both H3K23me1 and H3K23me2 were detected in the reaction product (Fig. 1B), although H3K23me3 was not detected. Future studies are needed to explain why our GST-SET-21 cannot produce H3K23me3 *in vitro* and determine whether SET-21 can produce H3K23me3 *in vivo*. Nevertheless, our results indicate that SET-21, the closest SET-32 homology in *C. elegans*, exhibits H3K23 HMT activity *in vitro*.

### SET-21 is expressed in *C. elegans* adult germline and embryo

Based on the published tissue-specific and developmental RNA-seq data sets ^41^ ^42^, both *set-21* and *set-32* mRNA expressions appeared to be germline-enriched (Fig. S1C and D). We performed immunofluorescence (IF) microscopy analysis using gonads dissected from adult hermaphrodite animals expressing the SET-21(native)::3xFLAG or SET-32(native)::3xFLAG protein. Our anti-FLAG IF microscopy of SET-32(native)::3xFLAG agrees with the previous study^36^ showing that SET-32 is expressed throughout the different developmental stages of adult germline (Fig. 2A, C). SET-32 is present in both the cytoplasm and nucleus.

**Figure 2.**
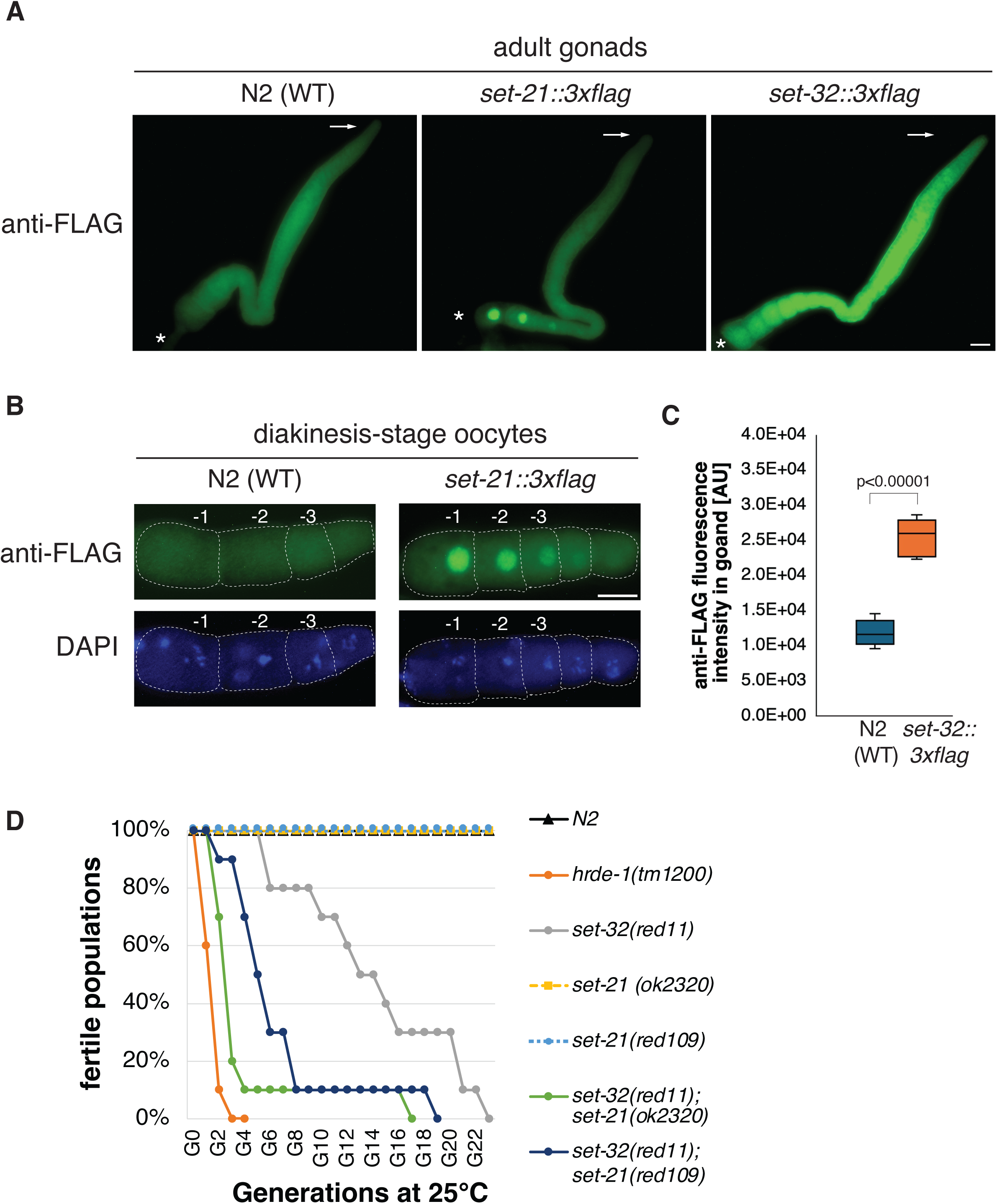
SET-21 and SET-32 are expressed in the adult germline and required for transgenerational fertility at an elevated temperature. (A) Representative anti-FLAG immunofluorescent (IF) images for dissected hermaphrodite adult gonads of N2 (WT), or animals expressing SET-21(native)::3xFLAG or SET-32(native)::3xFLAG. The distal and proximal tips of gonads were indicated with arrows and asterisks, respectively. Scale bar: 20μm. (B) anti-FLAG IF and DAPI images of diakinesis oocytes of WT and *set-21*(native)::3xFLAG animals. (C) Boxplot comparing anti-FLAG fluorescent intensity, measured by ImageJ in an arbitrary unit, between WT and *set-32*::3xFLAG gonads (N=5). The p-value is calculated by student’s *t*-test. (D) Transgenerational fertility assay was performed at 25°C. 10 lines were started for each strain and their progeny were transferred to a new plate at each generation until the population became sterile (See Methods for detail). The percentage of lines with fertile population was plotted as a function of the generation number for each strain. N2, *set-21(ok2320)*, and *set-21(red109)* exhibited 100% fertility throughout the assay.

We found that SET-21::3xFLAG was strongly enriched in the nuclei of oocyte diakinesis germ cells (Fig. 2A-B). Interestingly, the SET-21 expression progressively intensifies as diakinesis oocytes mature. Only a background level of SET-21::3xFLAG was observed in the earlier stages, including the mitotic proliferating and meiotic pachytene cells. We did not detect any expression of SET-21::3xFLAG in sperm. This result indicates that SET-21 expression is developmentally regulated and, in an adult animal, predominantly expressed in the oocyte nuclei.

We also performed anti-FLAG IF analyses in embryos. We found that both SET-21::3xFLAG and SET-32::3xFLAG proteins were broadly expressed in embryos (Fig. S2). Like adult germline, SET-32 is present in both the cytoplasm and nuclei of embryos, while SET-21 is strongly enriched in the nuclei of embryos.

### SET-21 and SET-32 promote germline immortality at a high temperature

To characterize the function of SET-21, we obtained a *set-21* mutant strain created by the genomic deletion consortium project ^43^. The *set-21(ok2320)* allele^43^ carries a 1.6 kb deletion that includes the entire catalytic SET domain of SET-21, likely resulting in a loss-of-function mutation. In addition, we constructed a putatively catalytic inactive mutation of *set-21* (Y502F, allele name *red109*). The tyrosine 502 residue is in the highly conserved Motif IV of the SET domain (Fig S1A) and its phenolic hydroxyl group is essential for the binding of S-adenosyl-methionine and catalysis in other HMTs ^38^. To examine any possible synthetic effect, we constructed two *set-32;set-21* double mutant strains, each carrying a different mutant *set-21* allele.

All of the *set-21, set-32,* and *set-32*;*set-21* mutants were continuously maintained in our 20°C incubator for at least one year without any sign of sterility. After shifting to 25°C, we found that the *set-32*;*set-21* double mutant animals, regardless which of the two aforementioned *set-21* mutant alleles was used, exhibited a progressive reduction in brood size and became sterile after approximated eight generations at 25°C (Fig. 2D and S3A-B). Such phenotype, termed mortal germline (Mrt), is common to the mutations in the germline nuclear RNAi pathway ^44, 45^. We examined *set-32;set-21* mutant animals cultured at 25°C for seven generations. We found that some mutant animals lacked sperm while some lacked oocytes or both types of gametes (Fig. S3C), indicating that both male and female germ cell development is defective in the mutant.

*set-32* mutant animals also exhibited the Mrt phenotype (Fig. 2D), consistent with the previous reports ^36, 46^ . Compared to the *set-32;set-21* mutant animals, it took much longer (>20 generations) for the *set-32* mutant to reach complete sterility in our analysis. Neither of the two *set-21* mutant strains showed any sign of the Mrt phenotype at 25°C (up to 23 generations). Our results indicated that SET-21 and SET-32 function synthetically to promote germline immortality at an elevated temperature.

### SET-32 and SET-21 are required for the H3K23me3 at the native nuclear RNAi targets

Knowing that SET-21 can methylate H3K23 *in vitro*, we decided to investigate the role of SET-21 in H3K23me *in vivo*. To this end, we performed H3K23me1, H3K23me2, and H3K23me3 ChIP-seq in the N2 (WT), *set-32*, *set-21*, and *set-32;set-21* young adult animals. To detect any obvious global changes, we made whole-chromosome coverage plots for each mutant in comparison with WT (Fig. S4). Each of the three mutants exhibited essentially WT-like profiles for all three H3K23me marks at the resolution we used for this analysis (10 kb).

We then increased the resolution to 1 kb for the mutant versus WT comparison (Fig. 3). We found that *set-21* mutation alone had virtually no effect on H3K23me1, me2, or me3 ChIP-seq signals. *set-32* single mutation had virtually no effect on H3K23me1 or H3K23me2, but resulted in modest decreases in H3K23me3 for 165 kb regions (cutoff: mutant/WT ≤ 2/3, FDR ≤ 0.05), which is consistent with our previous report ^21^. Mutating both *set-32* and *set-21* had no impact on H3K23me1, but showed modest decreases in H3K23me2, and significant losses in H3K23me3 (Fig. 3).

**Figure 3.**
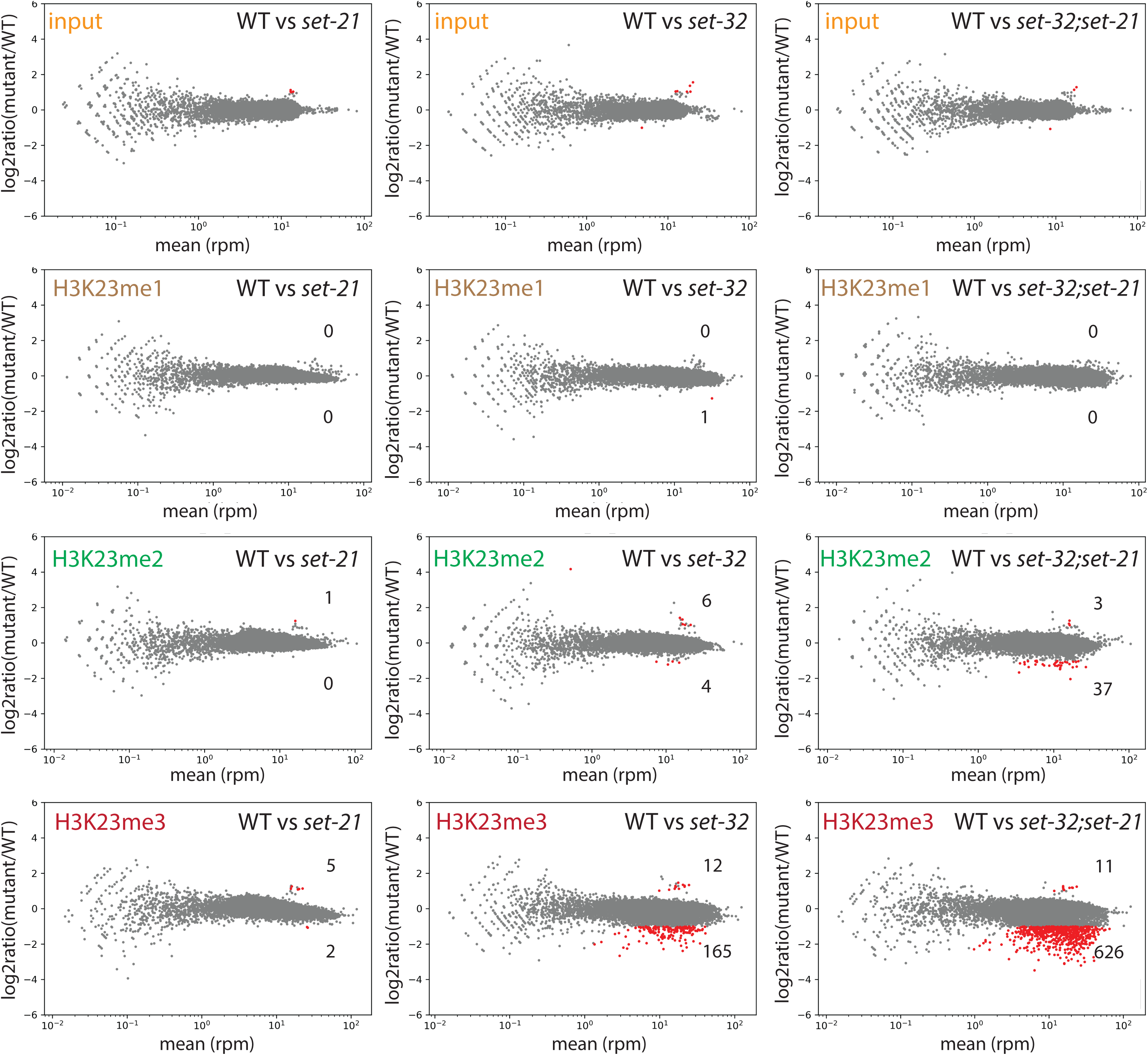
MA-plots of H3K23me1, me2, and me3 ChIP-seq comparing WT versus *set-21(red109)*, *set-32(red11)*, or *set-32(red11);set-21(red109)* mutant. Average RPM (reads per million sequenced tags) values from two replicates were calculated for each 1kb window throughout the whole genome. Regions with increased or decreased H3K23me in a mutant (highlighted in red) were determined by the BaySeq program ^74^ with a minimal 2-fold difference (FDR≤0.05), subtracting the regions that showed differential input signals (the top row). The numbers of regions with either increased or decreased H3K23me in a mutant were indicated in each panel.

To perform more detailed, quantitative analysis of H3K23me3 ChIP-seq data, we identified H3K23me3-enriched genomic regions using MACS2 ^47^. We first compared the H3K23me3 ChIP-seq and input signals in the WT animals, and identified 9918 H3K23me3 peaks, which covers approximately 5% of the genome (4.9 Mb) (Table 1). We then asked which of these peaks are dependent on HRDE-1 or SET-32/21 for H3K23me3. To this end, H3K23me3 ChIP-seq analysis was also performed for the *hrde-1* mutant in this study.

By comparing WT with *hrde-1* or *set-32;set-21* mutants, we identified 372 peaks (496 kb) and 408 peaks (512kb) in which the H3K23me3 enrichment is dependent on HRDE-1 or SET-32/21, respectively (cutoff: mutant/WT ≤ 2/3, FDR ≤ 0.05), (Fig. 3A and Table 1. See Fig. 5 for examples.). We found that the HRDE-1-dependent H3K23me3 peaks and the SET-32/21-dependent ones largely overlap (Fig. 4A), suggesting that SET-32/21-dependent H3K23me3 is largely limited to the germline nuclear RNAi targets.

**Figure 4.**
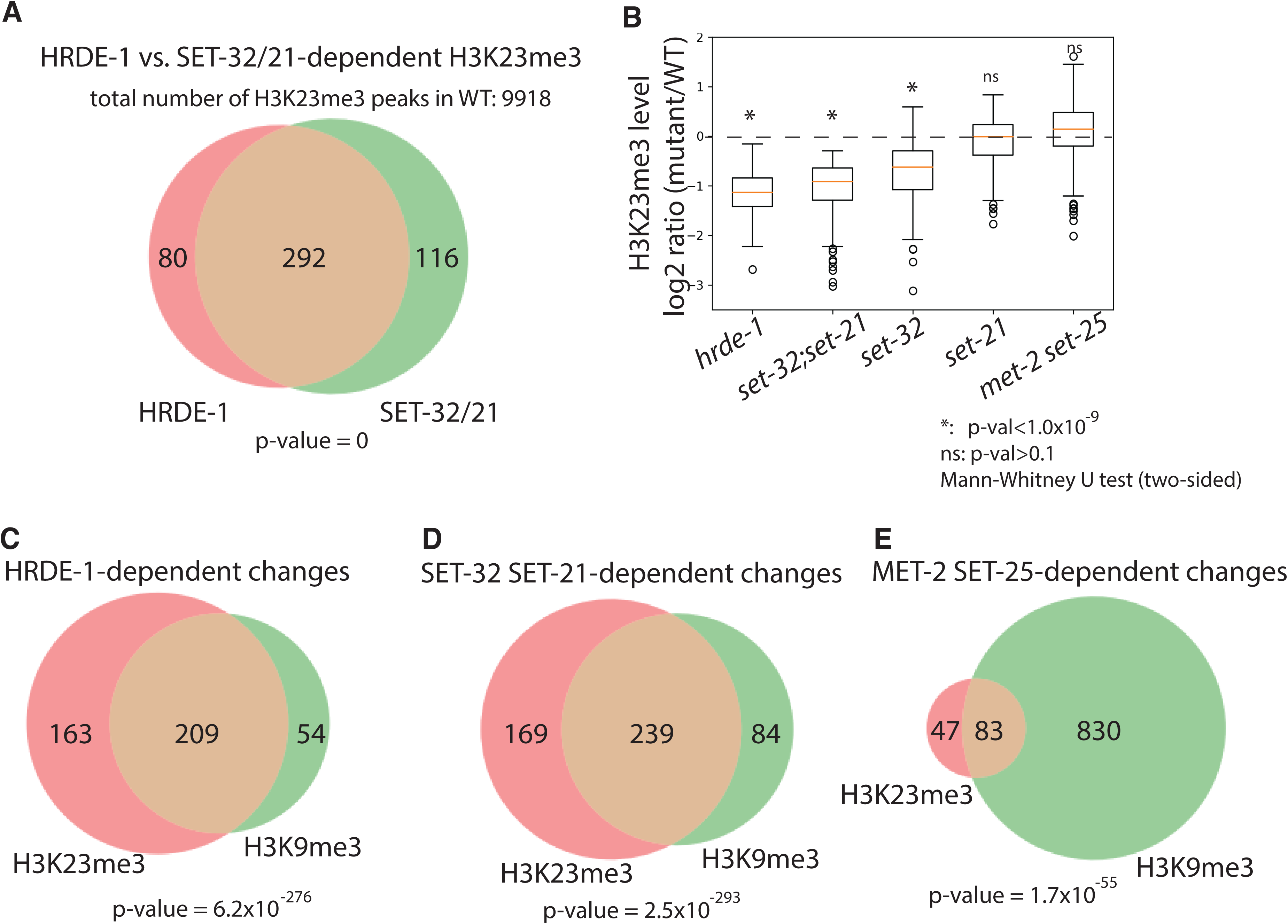
SET-32 and SET-21 are required for H3K23me3 and H3K9me3 at germline nuclear RNAi targets. (A) Venn diagram of numbers of regions with HRDE-1 and SET-32/21-dependent H3K23me3 (H3K23me3 [mutant/WT]≤2/3, FDR≤0.05). (B) Box plot of H3K23me3 levels (relative to WT) for regions with SET-32/21-dependent H3K23me3 in different mutant strains. Mann-Whitney U tests (two-sided) were used to determine the statistical significance for the H3K23me3 differences between a mutant and WT (null hypothesis: no difference). (C-D) Venn diagram of numbers of regions of H3K23me3 and H3K9me3 that are dependent on (C) HRDE-1, (D) SET-32 and SET-21, and (E) MET-2 and SET-25 (H3K23me3 or H3K9me3 [mutant/WT]≤2/3, FDR≤0.05). Hypergeometric distribution was used to calculate the p-values of the significance of the overlaps in the Venn diagrams (null hypothesis: no significant overlap).

**Figure 5.**
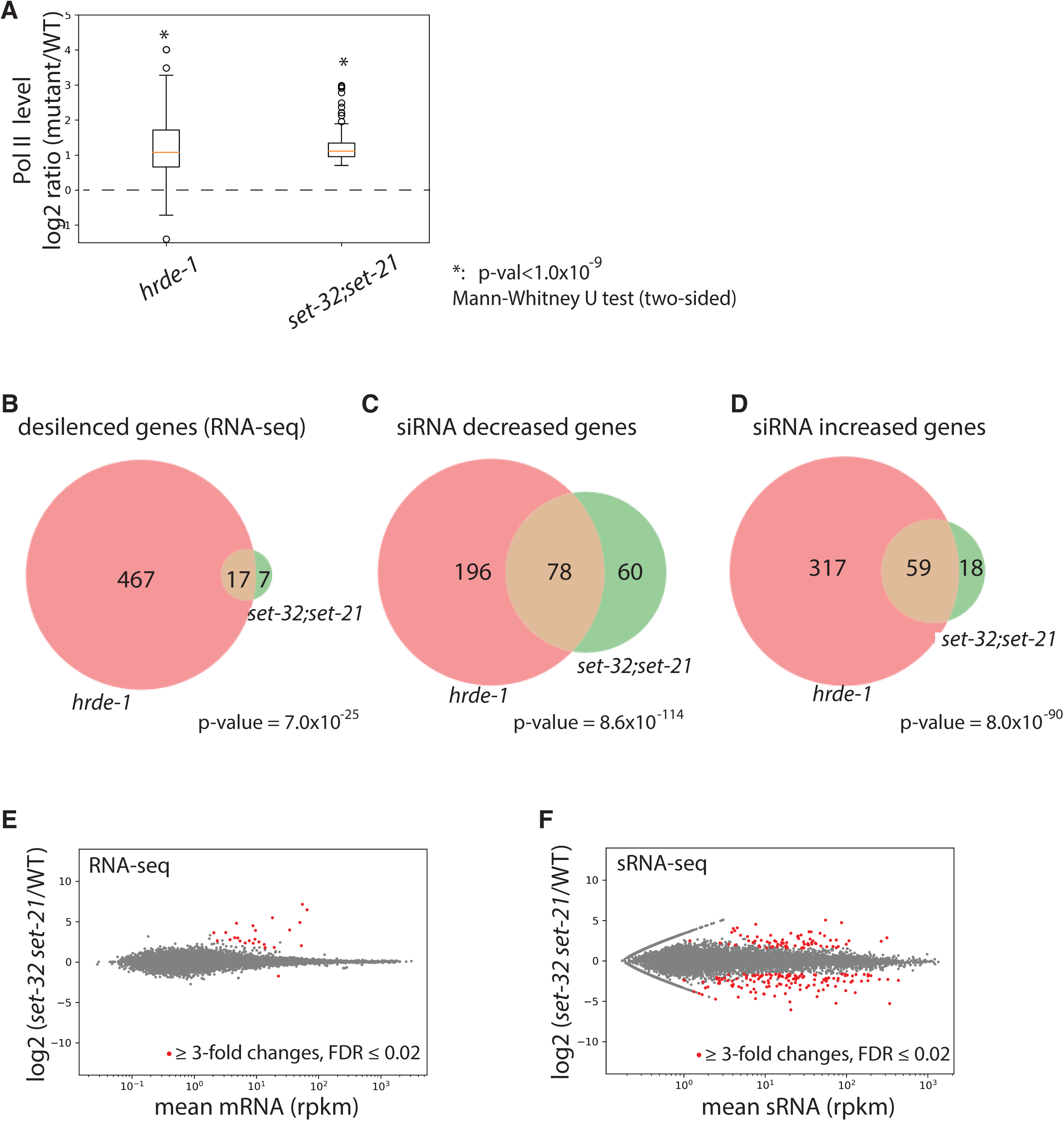
SET-32 and SET-21 are required for transcriptional repression and proper expression of siRNAs of germline nuclear RNAi targets. (A) Box plot of Pol II levels (relative to WT) for regions with SET-32/21-dependent H3K23me3 in *hrde-1* and *set-32;set-21* mutants. (B) Venn diagram of desilenced genes in *hrde-1* and *set-32;set-21* based on RNA-seq (cutoff: mutant/WT ≥ 3-fold, FDR ≤ 0.02). (C-D) Venn diagram of genes with decreased (C) or increased (D) siRNA expression in *hrde-1* or *set-32;set-21* mutant animals compared to WT (minimal 3-fold change, FDR ≤ 0.02). (E-F) MA-plots comparing WT and *set-32;set-21* mutant animals for (E) mRNA and (F) sRNA for all protein-coding genes.

Consistent with the 1kb whole-genome analyses (Fig. 3), *set-32* mutant exhibited weaker H3K23me3 loss in the SET-32/21-dependent regions than the *set-32;set-21* double mutant (Fig 4B and ^21^); while the *set-21* mutant did not show any obvious H3K23me3 losses in the same regions (Fig. 4B). We also performed H3K23me3 ChIP-seq in the *met-2 set-25* double mutant and found that loss of these two H3K9 HMTs did not cause significant reduction in the H3K23me3 level (Fig. 4B).

Our results indicate that SET-32 and SET-21 are germline nuclear RNAi-specific H3K23 methyltransferases. We note that, outside of the germline nuclear RNAi targets, most H3K23me3 peaks in *C. elegans* genome are independent of HRDE-1 or SET-32/21, indicating an RNAi-independent H3K23me3 pathway(s) and other unknown H3K23 HMTs.

### SET-32 and SET-21 are also required for the H3K9me3 at the native nuclear RNAi targets

SET-32 has been also shown to promote H3K9me3 *in vivo* in previous studies ^24, 36^. To investigate the possible role of SET-32/21 in whole-genome distribution of H3K9me3, we performed H3K9me3 ChIP-seq in WT and *set-32;set-21,* as well as *hrde-1* and *met-2 set-25* mutant animals.

H3K23me3 and H3K9me3 have almost the same genomic distribution in WT animals (Fig. S5A)^21, 33^. So we used the genomic annotations of the H3K23me3 peaks for the H3K9me3 analysis. We first determined the regions that showed at least 33.3% reduction (FDR ≤ 0.05) in H3K9me3 in *hrde-1*, *set-32;set-21*, or *met-2 set-25* mutant and called these regions with HRDE-1, SET-32 SET-21, or MET-2 SET-25-dependent H3K9me3, respectively.

We found that HRDE-1-dependent H3K9me3 and HRDE-1-dependent H3K23me3 are largely overlap (Fig. 4C). SET-32 SET-21-dependent H3K23me3 and SET-32 SET-21-dependent H3K9me3 also have very similar genomic distribution (Fig. 4D). In contrast, MET-2 SET-25-dependent H3K9me3 covers more genomic regions than MET-2 SET-25-dependent H3K23me3 (Fig. 4E and Fig. 6). In addition, the overlap between HRDE-1-dependent H3K23me3 (or H3K9me3) and MET-2 SET-25-dependent H3K23me3 (or H3K9me3) are much smaller than the ones between HRDE-1 and SET-32 SET-21 (Fig. S5B and C). These results suggest that germline nuclear RNAi-mediated H3K23me3 and H3K9me3 are two highly correlated events, and both are dependent on SET-32 and SET-21. Given the lack of the *in vitro* H3K9 HMT activity for SET-32 ^21^ and SET-21 (this study), we suggest that the H3K9me3 at the nuclear RNAi targets is deposited by MET-2 and SET-25 and is downstream to the activity of SET-32/21-mediated H3K23me3.

**Figure 6.**
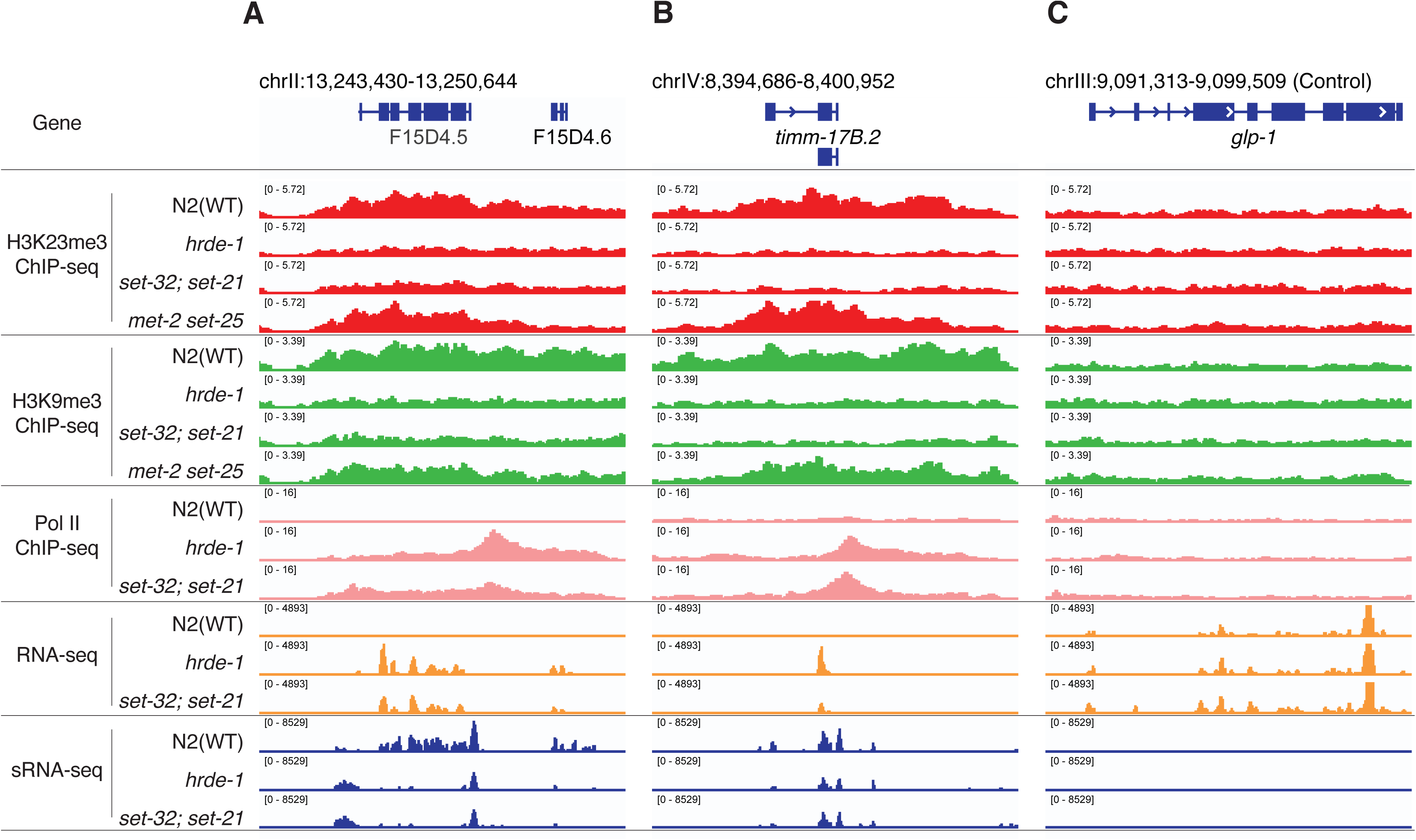
**Coverage plots of various ChIP-seq, RNA-seq, and sRNA-seq** for WT, *hrde-1, set-32;set-21, met-2 set-25* mutants at nuclear RNAi targets *f15d4.5* (A) and *timm-17b.2* (B), as well as a control euchromatin locus (*glp-1*) (C).

### SET-32 and SET-21 are required for transcriptional repression of a subset of germline nuclear RNAi native targets

We performed RNA-seq of WT, *set-32*, *set-21*, *set-32;set-21*, and *hrde-1* mutant animals. There are 484 protein-coding genes became derepressed in the *hrde-1* mutant using a minimal fold change [hrde-1/WT] of 3.0 (FDR ≤ 0.02) (Fig. S7A and Table 2), which is consistent with our previous studies ^22, 23^. Using the same cutoff, we found that only four and one genes were derepressed in the *set-32* and *set-21* single mutants, respectively. 24 genes were derepressed in the *set-32;set-21* mutant animals (Fig. S6A-B and Fig. 5E and Table 2). All four genes that were derepressed in the *set-32* mutant were also derepressed in the *set-32;set-21* double mutant. The single gene that was derepressed in the *set-21* mutant is likely due to genetic background because the same gene was not derepressed in *set-32;set-21* double mutant or a different set-21 allele (Fig. S6C). We named the 24 desilenced genes as *set-32/21*-sensitive targets. The siRNAs of *set-32/21*-sensitive targets are bound by HRDE-1 (Fig. S10A). 17 of the *set-32/21* sensitivity targets are also desilenced in the *hrde-1* mutant (Fig. 3B).

These results suggest that SET-32 and SET-21 play a redundant role in mRNA silencing. This is consistent with their redundant roles in H3K23me3 at nuclear RNAi targets and germline fertility. Majority of the 24 *set-32/21*-sensitive genes are germline nuclear RNAi targets. However, it is somewhat surprising that only a small fraction of the germline nuclear RNAi targets were desilenced in the *set-32;set-21* mutant despite that most of the germline nuclear RNAi targets showed loss of H3K23me3 in the *set-32;set-21* mutant.

To investigate the role of SET-32/21 in transcriptional repression, we performed Pol II ChIP-seq analysis of WT, *hrde-1*, and *set-32;set-21* mutant animals. We found that regions with SET-32/21-dependent H3K23me3 exhibited a strong tendency to have increased Pol II occupancies in both the *hrde-1* and *set-32;set-21* mutants (Fig. 5A and Fig. 6). Therefore, SET-32 and SET-21 are required for transcriptional repression at these native nuclear RNAi targets.

### Loss of SET-32/21 changes siRNA expressions for many genes

Previous studies have found intricate connection between chromatin enzymes and siRNA dynamics in *C. elegans* ^26, 48, 49^. To investigate the potential roles of SET-32/21 in siRNA regulation, we performed the sRNA-seq analysis. We found very few siRNA changes in the *set-21* mutant (Fig. S6E). In contrast, both *set-32* and *set-32;set-21* mutants exhibited extensive siRNA changes and the two mutants shared very similar siRNA profiles (Fig. S6D, S8C, S8D, and Table 2).

We found that the siRNA changes are more complex than the mRNA changes in the *set-32;set-21* mutant (Fig. 5E, 5F and Fig. S9). We found 138 genes had higher siRNA expression levels in *set-32;set-21* compared to WT animals and 77 genes had lower siRNA expression levels (cutoff: fold change ≥3.0 and FDR ≤0.02). Some of the top changes with increased mRNA expressions showed losses of siRNA expressions in the *set-32;set-21* mutant (Fig. 6 and S9A and S9C). Most of the siRNA changes, particularly for the genes with increased siRNA expressions, were not associated with any significant changes in mRNA expression in the *set-32;set-21* mutant (Fig. S9). Therefore, the impact of *set-32;set-21* mutations are far greater on the siRNA profiles than the mRNA profiles: affecting more genes and resulting both increased and decreased siRNA expressions.

Mutation in the key nuclear RNAi factor *hrde-1* also resulted in complex and extensive changes in siRNA expression profiles (Fig. S7B) ^22^. We found that the majority of siRNA changes in the *set-32;set-21* mutant also showed corresponding changes in the *hrde-1* mutant (Fig. 5C and 5D), indicating an overlapping role in siRNA regulation between HRDE-1 and SET-32/21.

We analyzed our published HRDE-1 and CSR-1 coIP sRNA-seq data^50^ and found that siRNAs showed differential expressions (either increase or decrease) in the *set-32;set-21* mutant tend to be bound by HRDE-1, instead of CSR-1 (Figure. S10), indicating that the *set-32;set-21* mutations selectively impact the WAGO-class secondary siRNAs (22G-RNAs).

### SET-32 and SET-21 only partially contribute to the transcriptional repression at germline nuclear RNAi targets

As mentioned in the previous section, some of the top *set-32/21*-sensitive targets (measured by mRNA changes) showed losses of siRNA expression in the *set-32;set-21* double mutant. We noticed that these genes also showed loss of siRNAs in the *hrde-1* mutant (*e.g. f15d4.5* Fig. 6A). Can restoring the siRNA expression rescue the transcriptional silencing at these targets in *hrde-1* or *set-32;set-21* mutant animals? To address this question, we used a piRNA-based gene silencing technology, termed piRNAi ^51^, which expresses a set of custom designed piRNAs from an extrachromosomal array. The ectopic piRNAs result in abundant secondary siRNAs against the target gene in the germline, which then leads to both classical and nuclear RNAi at the target gene ^51–54^.

Here we chose *f15d4.5* and *c38d9.2* as piRNAi targets. Both genes are annotated as putative protein-coding genes without any known functions in the Wormbase. They are native germline nuclear RNAi targets with abundant siRNAs in WT animals but exhibited loss of siRNAs and transcriptional derepression in the *hrde-1* and *set-32;set-21* mutants (Fig. 7B-D). We transformed the *hrde-1* and *set-32;set-21* mutant animals with a piRNAi transgene that expresses both anti-*f15d4.5* and anti-*c38d9.2* piRNAs. A control piRNAi transgene, which expresses a set of anti-randomly sequence piRNAs, was also introduced into the mutant strains. We first performed sRNA-seq and confirmed that the anti-*f15d4.5* and *c38d9.2* piRNAi, but not the control piRNAi, restored their siRNA expressions to the WT or even higher levels in both mutant strains (Fig. 7B).

**Figure 7.**
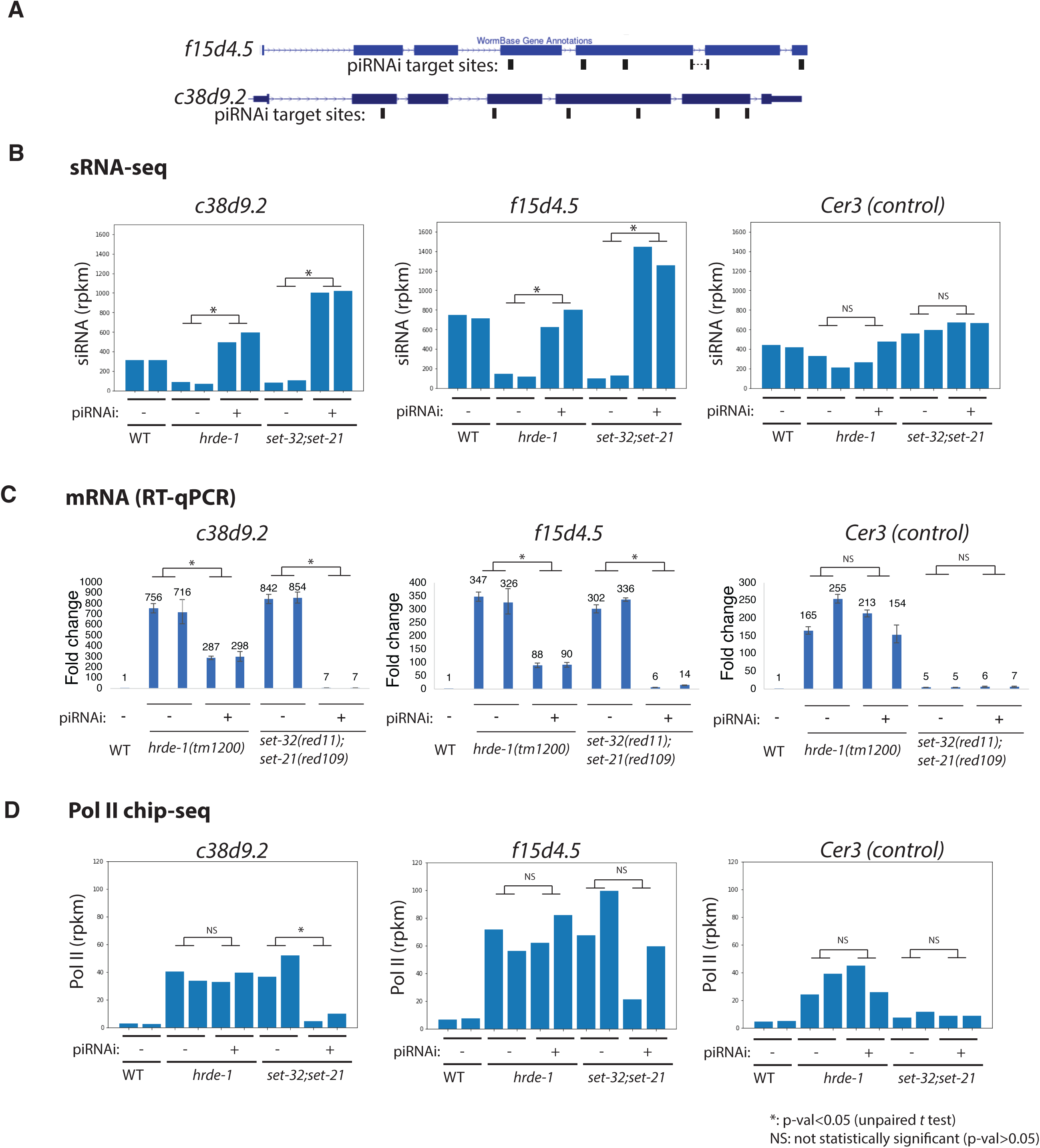
Transcriptional silencing defect at native targets of nuclear RNAi in *set-32;set-21* mutant can be partially rescued by piRNAi. (A) A piRNAi transgene targeting *f15d4.5* and *c38d9.2*, two native targets of germline nuclear RNAi, was introduced into *hrde-1* and *set-21;set-32* mutant animals. The piRNAi target sites are indicated in the schematic. (B-D) siRNA, mRNA, Pol II levels of *f15d4.5*, *c38d9.2*, and *Cer3* (an LTR retrotransposon not targeted by the piRNAi, as a control locus) in anti-f15d4.5+c38d9.2 piRNAi and anti-random control piRNAi, (labeled as piRNA + and -, respectively, in the figure) in WT, *hrde-1,* and *set-32;set-21* mutant backgrounds were shown. Two independent lines for each injection were used.

We then performed RT-qPCR and found that the anti-*f15d4.5*+*c38d9.2* piRNAi, but not the control piRNAi, was able to suppress their mRNA expressions in both *hrde-1* and *set-32;set-21* mutants (Fig. 7C). However, a much higher degree of suppression was observed in the *set-32;set-21* mutant than in the *hrde-1* mutant (*e.g.* 2.5-and 242-fold reductions in *c38d9.2* mRNA expressions in *hrde-1* and *set-32;set-21*, respectively). The partial rescue of silencing by piRNAi in the *hrde-1* mutant is consistent with the model that the piRNAi-induced secondary siRNAs rescue the PTGS, but not the TGS, as HRDE-1 is required for the TGS but not the PTGS mechanism. The near complete silencing by piRNAi observed in *set-32;set-21* mutant indicates that SET-32 and SET-21 are dispensable for silencing at these target genes when both HRDE-1 and siRNAs are present.

We performed Pol II ChIP-seq to investigate the role of SET-32/21 in the transcriptional repression when siRNAs are restored. First we observed that that anti-*f15d4.5* and *c38d9.2* piRNAi did not reduce the Pol II level at the target genes in the *hrde-1* mutant animals, which is consistent with the essential role of HRDE-1 in TGS. In *set-32;set-21* mutant, anti-*f15d4.5* and *c38d9.2* piRNAi reduced the Pol II level at these target genes by 83.8% and 50%, respectively, compared to control piRNAi in the same mutant. However, restoring the siRNAs in the *set-32;set-21* mutant did not fully rescue the transcriptional repression defect. The Pol II levels at *c38d9.2* and *f15d4.5* in the *set-32;set-21* (piRNAi+) were still 1.7 and 4.9 times higher than their WT levels (*i.e.*, when fully suppressed, Fig. 7D). The Pol II ChIP-seq results are consistent with mRNA-seq results, which showed a slight above the WT-level of mRNA expression of the two targets in the *set-32;set-21* mutant (piRNAi+).

Based on these results, we suggest that SET-32/21-mediated H3K23me3 is a partial contributor to the siRNA-guided transcriptional repression. For most of the native RNAi targets, loss of H3K23me3 can be compensated by other siRNA-guided TGS mechanisms, and therefore does not lead to significant desilencing at the mRNA level. The requirement of SET-32 and SET-21 for certain sensitive targets such as *f15d4.5* and *c38d9.2* may be due to siRNA loss or other unknown impacts of heterochromatin defects.

### The requirement of SET-21 and SET-32 for gene silencing triggered by exogenous dsRNA, piRNA, and transgene

Nuclear RNAi against a germline-expressed euchromatin gene can be induced by different exogenous trigger molecules, including dsRNA^8^, piRNA^52^, and extrachromosomal array^25^. Here we investigated whether SET-32 and SET-21 are required for these silencing pathways.

#### dsRNA

We fed WT, *hrde-1*, *met-2 set25*, and *set-32;set-21* animals with *oma-1* dsRNA-expressing *E. coli*. Two different *set-32;set-21* mutant strains were used, each carrying a different *set-21* mutant allele. RT-qPCR analysis of *oma-1* pre-mRNA indicated that the dsRNA feeding induced a strong transcriptional repression of *oma-1* in WT, *met-2 set25*, and *set-32;set-21* mutant animals (Fig. 8B). The *hrde-1* mutant was defective in the dsRNA-induced transcriptional repression at *oma-1* as previously reported^24, 44^. These results indicate that SET-32 and SET-21 are not required for dsRNA-induced transcriptional repression.

**Figure 8.**
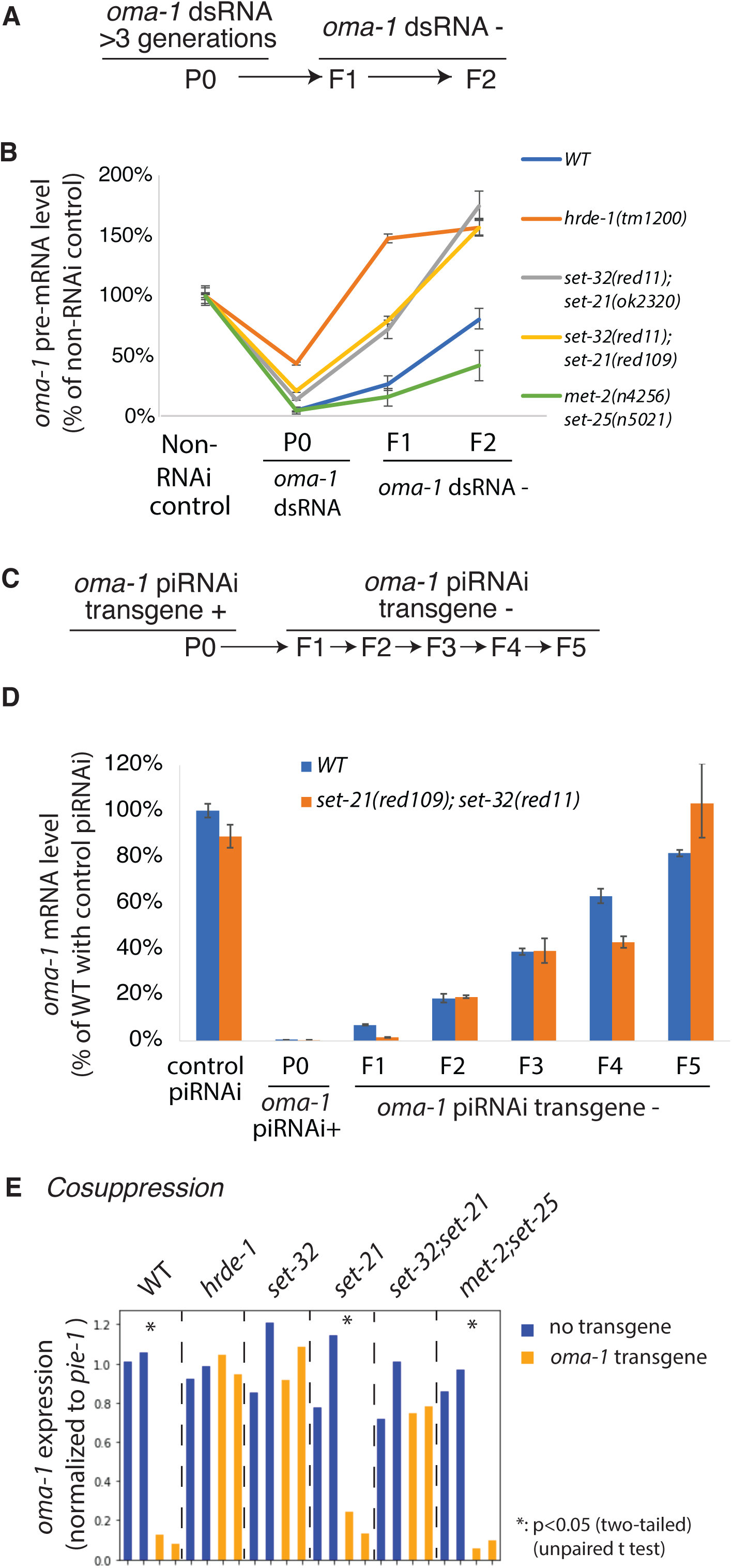
The requirement of SET-32 and SET-21 for (A,B) RNAi, (C,D) piRNAi, and (E) cosuppression. (A and C) The schematics of heritable RNAi by dsRNA feeding (A) and heritable piRNAi (C), both against *oma-1*. (B) *oma-1* pre-mRNA levels measured by RT-qPCR for heritable RNAi in WT and various mutants. (D) *oma-1* mRNA levels measured by RT-qPCR for heritable piRNAi. The values were normalized to the tubulin gene *tba-1* mRNA expression from the same sample and relative to the control samples. (E) mRNA levels of the native *oma-1* gene, measured by RNA-seq, for the cosuppression experiment in strains carrying the *oma-1* transgene. The values were normalized to a germline expressed gene *pie-1* mRNA levels.

**Figure 9.**
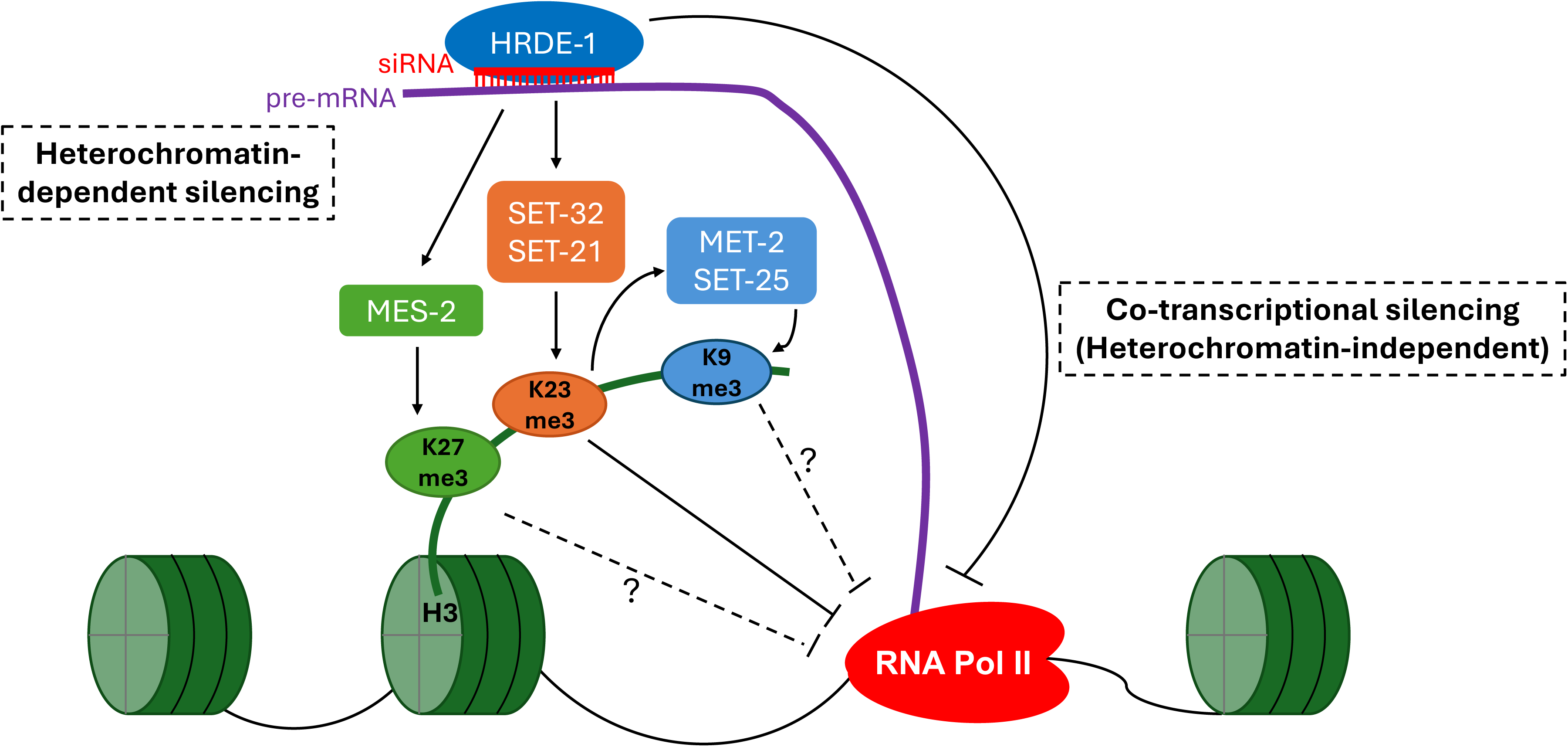
A model of germline nuclear RNAi-mediated heterochromatin pathway.

To measure the heritable RNAi effect, we collected two generations of progeny (F1 and F2) after dsRNA feeding had been discontinued (Fig. 8A). *Oma-1* silencing, measured by the pre-mRNA level, persisted in both F1 and F2 generations in the WT animals, but completely dissipated in the F1 generation for the *hrde-1* mutant (Fig. 8B) as expected ^24, 44, 52^. *met-2 set-25* showed an enhanced heritable RNAi compared to WT, which is likely due to the antagonistic role of MET-2 in heritable RNAi as previously reported ^26^. In both *set-32;set-21* mutant strains, *oma-1* silencing occurred in the F1 generation, but the degree of silencing was weaker than the WT animals (Fig. 8B). At the F2 generation, the heritable silencing effect was completely lost for the *set-32;set-21* mutants. These results indicate that SET-21 and SET-32 are required for a robust heritable RNAi effect induced by dsRNA.

#### piRNA

We performed piRNAi against *oma-1* in WT and *set-32;set-21* mutant animals. We examined the piRNAi transgene-containing animals, as well as the descendants that had lost the piRNAi transgene for one or several generations (Fig. 8C). This allowed us to examine the piRNA-induced heritable gene silencing effect. RT-qPCR was performed to measure the *oma-1* mRNA expression. We found that *oma-1* piRNAi silenced *oma-1* mRNA expression in both WT and *set-32;set-21* animals (Fig. 8D). The heritable silencing profiles shown in the transgene-negative descendants were also similar between WT and *set-32;set-21* mutant animals (Fig. 8D). These results indicate that SET-32 and SET-21 are not required for exogenous piRNA-induced silencing, either at the piRNA(+) generation or the heritable silencing.

#### Cosuppression

In addition to dsRNA and piRNA, a germline gene in *C. elegans* can also be heritably silenced by a homologous extrachromosomal transgene, a phenomenon called cosuppression ^55–57^. DNA fragment injected into *C. elegans* syncytial germline forms DNA repeat structure called extrachromosomal array ^58^. The repetitive DNA nature of such transgene has been suggested as the triggering signal for silencing both the transgene and the homologous native gene in the germline ^59^. Cosuppression shares some of the mechanisms of RNAi, including secondary siRNAs and heritable silencing ^25, 55, 56^. Much of its mechanism, especially the early steps of the pathway, is not well understood.

We transformed worms with an extrachromosomal transgene array carrying a partial *oma-1* cDNA driven by the *oma-1* promoter. The 492 nt *oma-1* cDNA fragment covers exons 2-4 and contains a SNP every 30 nt. We found the *oma-1* transgene caused a 10-fold reduction in *oma-1* mRNA expression in WT animals (Fig. 8E). We also observed strong cosuppression effect in *met-2 set-25 double* (11-fold) and *set-21* mutant animals (5-fold). The cosuppression effect was defective in *set-32, set-32;set-21,* and *hrde-1* mutant animals. These results indicate that the germline nuclear RNAi pathway and H3K23me3 is essential for cosuppression.

## Discussion

In this study, we identified a novel H3K23 HMT, SET-21. Together with SET-32, these two HMTs deposit most if not all H3K23me3 specifically at the germline nuclear RNAi targets, and function in synergy to promote transgenerational RNAi and fertility. Our work deepens the understanding of nuclear RNAi, especially the complexity of chromatin regulation and its connection to transgenerational epigenetic inheritance.

### The relationship between SET-32 and SET-21

SET-21 is the closest homolog of SET-32 in *C. elegans*. The two genes also have similar gene structures in terms of exon and intron organization (Fig. S1B), suggesting that they are likely to be evolved from a gene duplication event. However, the two genes are not completely redundant of each other, indicated by stronger phenotypes (e.g., H3K23me3 loss and Mrt) shown by the *set-32* mutant than the *set-21* mutant. On the other hand, SET-32 alone is not sufficient to replace SET-21 evidenced by a much enhanced phenotype of the *set-32;set-21* double mutant compared to the *set-32* single mutant. It is possible that the apparent synergy between SET-21 and SET-32 is due to their differential expression within the germline tissue: SET-21 expression is limited to oocytes while SET-32 is expressed throughout the different stages of adult germline. It is also possible that SET-21 and SET-32 have different biochemical activities. Future studies are needed to investigate these hypotheses. *set-32* and *set-21* have also been reported to extend the life span of *daf-2* mutant animals. SET-32 and SET-21 appear to function in the same pathway instead of synergistically for the life span phenotype^60^, suggesting their role in life span may be independent of their nuclear RNAi function.

### SET-32/21-dependent H3K23me3 is specific to the germline nuclear RNAi pathway

Our ChIP-seq analysis showing that HRDE-1- and SET-32/21-dependent H3K23me3 has very similar genomic profiles, which account only for approximately 10% of all H3K23me3-enriched regions in the genome. This probably explains why we were not able to detect H3K23me3 loss in the *set-32;set-21* mutant by either western blotting or mass spectrometry (data not shown).

H3K23me3 in nuclear RNAi-independent heterochromatin is not affected in the *set-32;set-21* mutant, suggesting the existence of addition H3K23 HMTs. Identifying the unknown H3K23 HMTs will be important to investigate the broader function of H3K23me3, which is abundant in *C. elegans* ^34^.

We do not understand how SET-32 and SET-21 are recruited to the nuclear RNAi targets at this point. Previous proteomic studies did not detect SET-32 or SET-21 in the HRDE-1 co-immunoprecipitation experiment ^61^, suggesting that either HRDE-1 and SET-32/21 interactions are very weak, or SET-32/21 were indirectly recruited to chromatin targets by HRDE-1.

### The relationship between H3K23me3 and H3K9me3

A previous study reported that MET-2 and SET-25 are the sole H3K9 HMTs in the embryo^62^. We previously found that adult *met-2 set-25* double mutant had only a partial loss of H3K9me3 at nuclear RNAi targets ^24^. In the *met-2 set-25;set-32* triple mutant, the H3K9me3 level was reduced to the background level ^24^. A stronger loss of H3K9me3 was also reported in *set-25;set-32* mutant germline compared with *set-25* or *set-32* single mutant ^36^. Interestingly, we also observed strong loss of H3K9me3 at germline nuclear RNAi targets in the *set-32;set-21* mutant in this study. Based on these results, we suggest a model in which MET-2 and SET-25 only partially contribute to RNAi-dependent H3K9me3, and additional H3K9 HMTs also function in the nuclear RNAi pathway.

SET-32/21-dependent H3K9me3 and H3K23me3 profiles correlate well with each other. It is conceivable that the both H3K9me3 and H3K23me3 at nuclear RNAi targets are deposited by SET-21 and SET-32, as mammalian EHMT1/GLP and EHMT2/G9a are known to deposit both H3K9me and H3K23me ^63–65^. Our HMT assays argue against this possibility. However, we cannot rule out that SET-32, SET-21, or both can deposit both H3K23me3 and H3K9me3 *in vivo*. It is also possible that an unknown HMT functions in a H3K23me3-dependent manner to deposit H3K9me3 at the nuclear RNAi targets. Future study is needed to test these hypotheses.

### The transcriptionally repressive role of H3K23me3

Our Pol II chip-seq analyses indicate that SET-32 and SET-21 promote transcriptional repression at germline nuclear RNAi targets. However, both H3K9me3 and H3K23me3 levels are reduced in the *set-32;set-21* mutant. Which of the two histone modifications contribute to transcriptional repressive? We currently favor H3K23me3 for this role because (1) H3K23me3 is much more abundant than H3K9me3 in *C. elegans* ^34^, and (2) the near complete loss of H3K9me3 in the *met-2 set-25;set-32* triple mutant did not exhibit transcriptional repression defect. However, we cannot rule out both H3K23me3 and H3K9me3 are needed for the transcriptional repression. We note that not all nuclear RNAi targets that showed increased Pol II occupancy in the *set-32;set-21* mutant. The *hrde-1* mutant had more desilencing events, measured by mRNA levels, than the *set-32;set-21* mutant. Based on these results, we suggest the following model: SET-32/21-dependent H3K23me3 repress the chromatin access of Pol II. But this is not the sole silencing mechanism of nuclear RNAi. Other HRDE-1-guided activities may eliminate the transcripts through an unknown co-transcriptional silencing mechanism.

### SET-32 and SET-21 as regulators of siRNA homeostasis

Interestingly, mutations in *set-32* and *set-21* led to a much more extensive changes in global siRNA expression pattern than mRNA. We note that some genes showed decreased siRNA expression, while some other genes showed increased siRNA expression. This suggests that the impact of SET-32 and SET-21 on siRNA expression is likely to be indirect. In some targets, the loss of siRNAs in the *set-32;set-21* mutant can potentially explain their silencing defects, evidenced by partial rescue of transcriptional silencing defect in the *set-32;set-21* mutant by re-introducing corresponding siRNAs.

### SET-32 and SET-21 are required for TEI in gene silencing and transgenerational fertility

Previous studies showed that SET-32 promotes the establishment of transgenerational epigenetic silencing either at some native germline nuclear RNAi targets or exogenous dsRNA-induced heritable RNAi ^17, 48^. Here we showed that SET-32 and HRDE-1 are also essential for transgene-induced silencing (cosuppression) in *C. elegans* germline. Furthermore, SET-32 and SET-21 function together to promote exogenous dsRNA-induced heritable RNAi. Similar to other germline nuclear RNAi factors, loss of SET-32 and SET-21 leads to the mortal germline phenotype at an elevated temperature. These results indicate that SET-32 and SET-21 are key TEI factors in *C. elegans*. Further investigation of the molecular and developmental mechanisms of these two enzymes and H3K23me3 should provide insight of novel aspects of TEI in animals.

## Methods

### Worm strains

*C. elegans* strain N2 (PD1074) is a cloned population derived from the original “Bristol” variant of *C. elegans* ^66^ and was used as the standard WT strain. Alleles used in this study were LG I: *set-32(red11)*, LG III: *hrde-1(tm1200), met-2(n4256) set-25(n5021),* LG IV: *set-21(ok2320), set-21(red109).* N2(PD1074), *hrde-1(tm1200), met-2(n4256), set-25(n5021),* and *set-21(ok2320)* strains were acquired from *Caenorhabditis* Genetics Center (CGC). We constructed the s*et-21(ok2320);set-32(red11)* or *set-21(red109);set-32(red11)* double mutant by CRISPR method as described in^67, 68^. *C. elegans* culture was as previously described^69^ in a temperature-controlled incubator. Worms were cultured at 20°C for all experiments except the multigenerational fertility assay at 25°C.

**Phylogenetic analysis of the 38 C. elegans SET-domain containing proteins** were performed using the Clustal Omega program ^70^ with the default setting.

### GST-SET-21 protein purification

SET-21 cDNA was prepared by RT-PCR using *C. elegans* mRNA and cloned into the pGEX-p6-1 vector. GST-SET-21 protein was obtained using a protein expression and purification procedure previously described in ^21^.

### *In vitro* HMT assay and mass spectrometry

The 75 µL HMT assay mixture contained 0.15 µM GST-SET-21, 213 µM S-adenosylmethionine, 2.5 µM H3.1, in 1X HMT buffer (50 mM Tris-HCl, pH 8.0, 20 mM KCl, 10 mM MgCl2, 0.02% Triton X-100, 1 mM DTT, 5% glycerol, and 1 mM PMSF). The reaction was incubated for 2 hours at 20°C. The histone peptides were prepared and analyzed by mass spectrometry as described in ^21, 71^.

### Brood size analysis and multigenerational fertility assay

Multigenerational brood size analysis was performed at 25°C (restrictive temperature). Worms were maintained at 20°C (permissive temperature) without starvation for at least 5-6 generations before the brood size analysis. Ten L2/L3 worms from each strain were transferred to one plate at 25°C. Their progeny, which grew at 25°C since 1-cell embryo, was considered as F1. When the F1 animals reached the L4 stage, ten worms from each strain were transferred to a new plate as the maintenance plate.

Another ten worms were individually placed onto new plates to count their brood size. These worms were transferred to a new plate each day during their egg-laying stage to facilitate counting. This procedure was repeated until at which *set-21;set-32* double mutant animals became completely sterile. A single-generation brood size analysis was performed for 20°C.

Germline fertility assay was performed at 25°C. At the first generation, 5 worms from 20°C were transferred to a new plate, which was then incubated at 25°C. 10 plates total for each strain were started as 10 lines. After three days, if less than 5 progenies were observed, we consider the line to be terminated. Otherwise, the line is considered viable and 5 progenies were transferred to a new plate. At each generation, the percentage of viable lines were calculated and used to generate the survival plot.

### Immunofluorescence

Adult worm gonads were dissected and fixed in 3% PFA in 100mM K2HPO4 for 5min and were wash three times in PBST (1xPBS with 0.1% Tween-20). Then the samples were permeabilized in 100% methanol at –20°C for 5 mins, washed in PBST for three times, and blocked in 0.5%BSA in PBST for 30 minutes at room temperature. The gonads were incubated in 1:200 mouse-anti-FLAG (Sigma) primary antibodies for two hours at room temperature, washed three times in 0.5% BSA in PBST, and then incubated in 1: 200 Donkey-anti-Mouse IgG-Alexa 488 (Jackson ImmunoResearch Laboratories) for one hour at room temperature. After three washes in 0.5% BSA in PBST for ten minutes, the gonad was mounted to 2% agar pad for imaging.

For embryo immunofluorescence staining, synchronized young adult worms (24 hours post-L4 stage, 20°C) were dissected in water on a poly-L-Lysine slide to release embryos. The slides were snap frozen in liquid nitrogen with coverslip on, and then were immediately incubated in -20°C methanol for 5 minutes after the coverslip was popped off the slide. After washing the slides three times in PBST (1xPBS with 0.1% Tween-20), the embryos on slides were blocked in blocking solution (0.5%BSA in PBST) for 20 minutes, incubated in 1:100 mouse anti-FLAG antibody (Sigma) for 1 hours, washed three times in PBST, and then incubated in 1:100 anti-mouse-IgG Alexa-488 (Jackson ImmunoResearch Laboratories) for 30 minutes. Slides were then stained with DAPI, washed three times in PBST, mounted with Slowfade (ThermoFisher Scientific) for imaging.

Fluorescence images were obtained using an Epi-fluorescence microscope: Zeiss Oberver.Z1 microscope equipped with ORCA-Flash4.0 LT Digital CMOS camera (Hamamatsu) and oil-immersion objective (40x). Images were captured using Metamorph 7.10. as a 16-bit single-plane image (For gonads: exposure time 8000ms for Alexa-488, and 4000ms for DAPI, without saturating pixels. For embryos: exposure time 4000ms for Alexa-488, and 500ms for DAPI, without saturating pixels.).

Gonad fluorescence was quantified using Fiji (ImageJ). Only intact gonads were used for measurement. Whole gonad area was manually selected, then the florescence level was measured using he Analyze->Measure function of Fiji. The mean grey-scale value in the measurement result was used for statistical calculation in Fig. 2C. The images in Fig. 2 and Fig. S2 were presented using identical brightness and contrast.

### piRNAi

The piRNAi transgene fragments were designed by using the wormbuilder webtool (www.wormbuilder.org/piRNAi) according to ^51^. The piRNAi target sites of piRNAi against C38D9.2 and F15D4.5 were illustrated in Fig. 7. The piRNA sequences targeting C38D9.2 are 5’-UCACAGGAGAUUCCUUUCGUG-3’, UCGGUGAGGAUUGAUUGGAAU, UCAGGAGGUUUGGUGUAAUCU, UCCGGUAAGUUUUUGCACAGC, UGGGCAGUUGGUAUGCAUUUG, and UCGGACGUUCUUGGGUAUUAU. The piRNA sequences targeting F15D4.5 are UCCGUUUCGCUUGCUGCGUUG, UGAGAGUUUGUCGUCUACCUU, UGGGCUUGUUCGACGCGGUUG, UAGCUUCUGCCAAGGUGGAAU, UGCAGGUAUUCUCGACUCCCU, and UGACGUCCUCCUCUGUUGGAA. anti-oma-1 piRNAi fragment was designed by ^51^. piRNAi DNA fragments were ordered from Twist DNA. The piRNAi transgenic animals were constructed according to ^51^.

### Cosuppression

Worms were injected with 60 ng/ µL pSG32 (oma-1 suppression plasmid), 20 ng/ µL pPD93_97 (myo-3p:GFP), 20 ng/ µL IR98 (Hygromycin resistance). pSG32 was constructed by inserting a transgene fragment into the pCFJ350 vector. The transgene is driven by the *oma-1* promoter and includes a 492 nt partial *oma-1* cDNA fused with a 1646 nt partial *smg-1* genomic DNA including the last five exons and intervening introns. The *oma-1* cDNA fragment covers exons 2, 3, and 4 and contains SNPs every 30 nt to distinguish from the native WT *oma-1*. Transgene-carrying worms were selected by hygromycin and confirmed by GFP expression. Two independent transgenic lines were used for each genetic background. Synchronized young adult animals were used for RNA-seq analysis.

### Preparation of worm grinds

Preparation of worm grinds has been described in ^22^. Briefly, synchronized L1 worms were prepared using the hypochlorite bleaching method, and then were released on NGM containing E. coli OP50. The synchronized worms reached young adult stage after 68 hours at 20°C and were harvested by washing off the plates by M9 buffer. Bacteria were removed by centrifugation of worms in a clinical centrifuge in a M9 buffer with 10% sucrose. Worms were then pulverized by grinding in liquid nitrogen with a mortar and pestle and were stored at −80°C.

### RT-qPCR

Total RNA was extracted from adult worm grind using Trizol reagents (ThermoFisher) according to manufacturer’s instructions. Total RNA was treated with DNase I (NEB) followed by phenol chloroform extraction. Then cDNA synthesis was performed using Reverse transcriptase III (ThermoFisher) as described in the manufacture’s manual. Quantitative PCR was performed using a QuantiStuido 3 real time PCR system using SYBR green master mix (ThermoFisher). ΔΔCT method was used to calculate the relative transcript abundance. *tba-1* was used as endogenous control.

### RNA-seq library preparation

Total RNA was first extracted from adult worm grind using Trizol reagents (ThermoFisher), then ribosomal RNA (rRNA) was depleted using RNaseH and PAGE-purified DNA oligos that are antisense to rRNA as described previously^72^. The rRNA-removed RNA was used to construct barcoded RNA-seq libraries using the SMARTer Stranded RNA-Seq Kit (Takara).

### sRNA-seq library preparation

Small RNA was extracted using mirVana miRNA isolation kit (ThermoFisher). The small RNA libraries were constructed using a 5’ - monophosphate independent, 3’ and 5’ linker ligation-based methods as previously described ^22^. The stranded Hi-seq index was added to the primer at the PCR steps to allow multiplexing.

### ChIP-seq library preparation

Chromatin immunoprecipitations were performed using the protocol described in ^21^. Briefly, grind of approximately 5000 adult worms was corsslinked and then sonicated to 200-500bp. Immunoprecipitation was performed using the following antibodies: anti-H3K23me3 (61500, Active Motif), anti-H3K9me3 (ab8898, Abcam). The pulled-down complexes were reversed crosslinked by proteinase K digestion and then purified by phenol chloroform extraction. The yielded DNA was used to construct barcoded ChIP-seq libraries using the KAPA Hyper Pre Kit (Roche) according to the manufacturer’s instruction.

### High-throughput sequencing

Uniquely barcoded RNA-seq, sRNA-seq, and ChIP-seq libraries were pooled and then sequenced on the Illumina HiSeq or Illumina NovaSeq X Plus instrument. Library names and list are in supplemental table (Table 3).

## Bioinformatic analysis

### ChIP-seq data analysis

#### H3K23me3 peak calling

Regions enriched for H3K23me3 in WT animals were determined for each of two sets of WT H3K23me3 ChIP-seq experiments by using macs2 ^47^. The command line is “macs2 callpeak -t [ChIP bam file] -c [ChIP input bam file] -g ce --outdir [output folder] -n [experiment name] - -nomodel --extsize 147 -m 5 100 -q 0.1 --broad”. The overlapping peaks from the two different WT experiments were identified using the “bedtools intersect” program ^73^. We did not merge nearby peaks because such merge reduces the sensitivity of calling differential regions.

#### Differential analysis of ChIP-seq

Differential H3K23me3 between WT and mutant animals were determined by using the BaySeq program^74^. Only the H3K23me3 peaks in the WT animals, as determined in the previous section, were used for this analysis. Two sets of experiments were performed for both WT and the mutant strain. Differential regions were separately determined for the H3K23me3 ChIP libraries and input libraries. Fold of change >= 1.5 and FDR <=0.05 were used as cutoff to determine the differential regions. Differential regions that were found in both the input libraries and ChIP libraries were removed from the final list.

#### RNA-seq analysis

For RNA-seq libraries prepared with the SMARTer Stranded RNA-Seq Kit (Takara), 51 nt segment of the R1 reads were used for sequence alignment against the mRNA sequences of C. elegans protein-coding genes using bowtie 1.2.3 ^75^. For RNA-seq libraries prepared with the 3’-linker ligation method, R1 reads with 5’-barcodes (4 nt) and 3’-linker sequence removed were used for the alignment. The number of perfectly aligned reads for protein-coding genes were used to determine the differentially expressed genes by using the BaySeq software ^74^ with default parameters. sRNA-seq libraries were similarly analyzed as RNA-seq libraries except that only 20-24 nt reads that were antisense to the mRNA sequences were used.

#### Venn diagram, boxplot, MA plots, were generated using python

Data availability: High-throughput sequencing data associated with this study has been deposited in NCBI GEO database with accession numbers of GSE266182, GSE266183, and GSE266184. Mass spectrometry data has been deposited in ProteomeXchange (https://www.proteomexchange.org/) with accession number of PXD052034 (Username; Password: nvXjzFdr).

## Acknowledgement

We thank Helen Ushakov and Elaine Gavin for technical assistance. Some strains were provided by the CGC, which is funded by NIH Office of Research Infrastructure Programs (P40 OD010440). Research reported in this publication was supported by the Busch Biomedical Grant from Rutgers, The State University of New Jersey, the National Institute of General Medical Sciences of the National Institutes of Health (R01GM111752 and R35GM152219) to S.G.; Research Foundation Flanders – FWO for personal funding (1SF2622N) and awarding a mobility grant (V400623N) to L.C.; the Hevolution Foundation (AFAR) award, the Einstein-Mount Sinai Diabetes center, and the NIH Office of the Director award (S10OD030286) to S.S.

---

**Figure S1.**
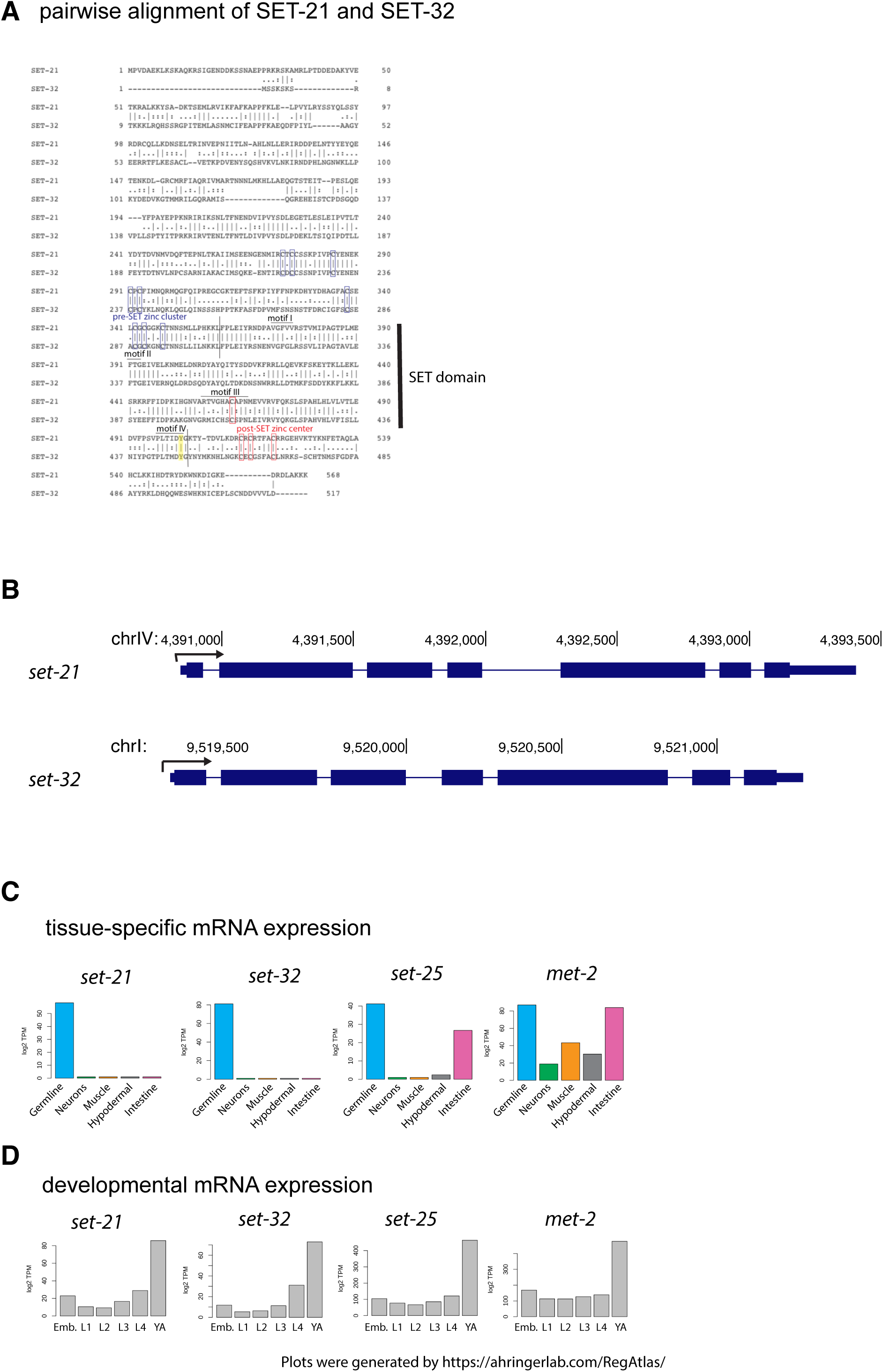
(A) Pairwise alignment of SET-21 and SET-32 proteins. Motifs I-IV of the SET domain, pre-SET zinc cluster, and post-SET zinc center were highlighted. The SET domain was marked by vertical lines. (B) Genome browser shots of *set-21* and *set-32* genes. (C-D) Tissue-specific and developmental mRNA expression profiles for *set-21, set-32, set-25*, and *met-2* using data generated by ^41, 42^. Plots were generated by https://ahringerlab.com/RegAtlas/.

**Figure S2.**
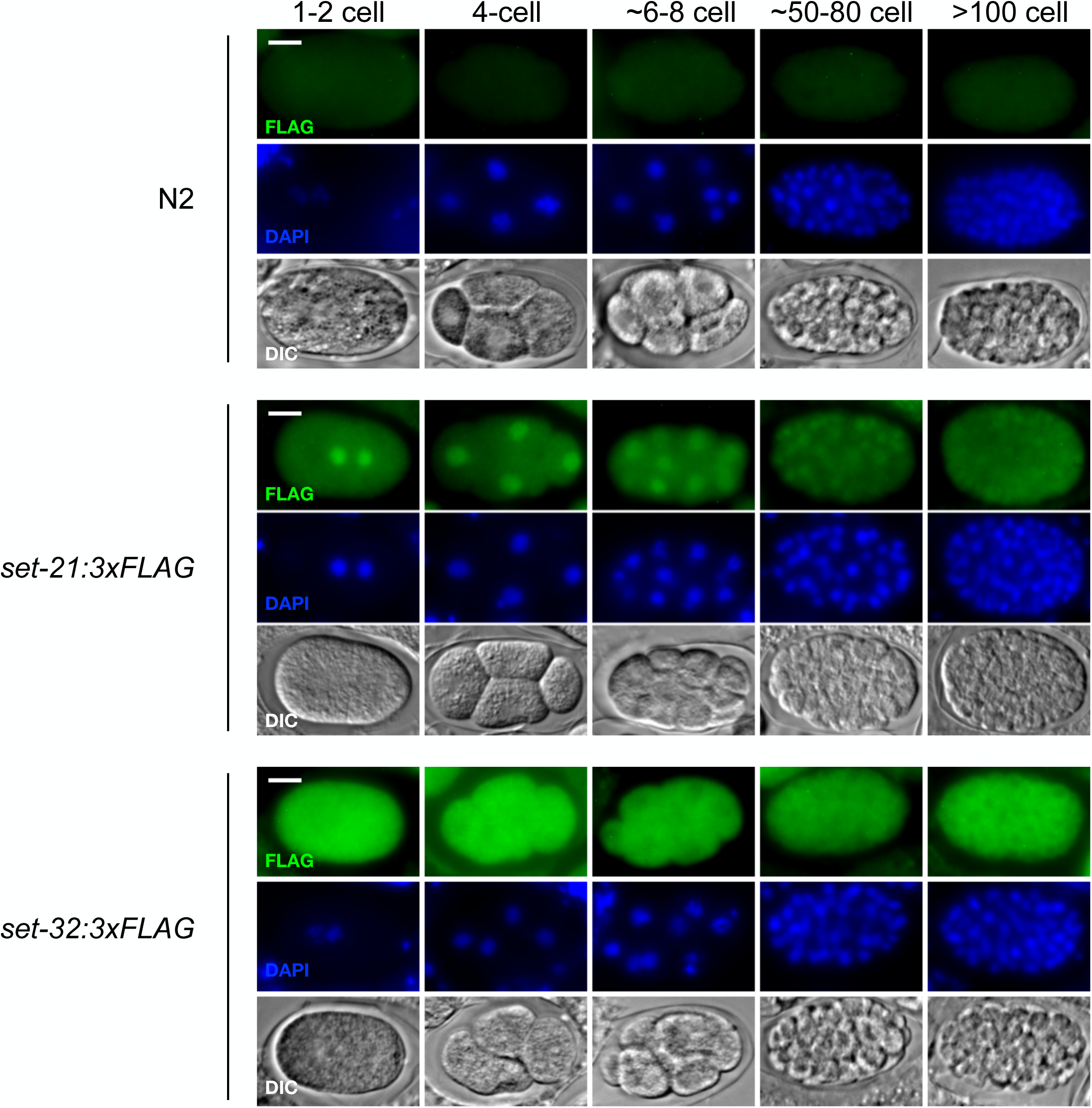
SET-21 and SET-32 are expressed in embryo. Anti-FLAG immunofluorescent microscopy was performed for different stages of N2, *set-21*(native)::3xFLAG, *set-32*(native)::3xFLAG embryos. Representative IF images were showed for each strain, together with DAPI and DIC images of the same embryo. Scale bar: 10μm.

**Figure S3.**
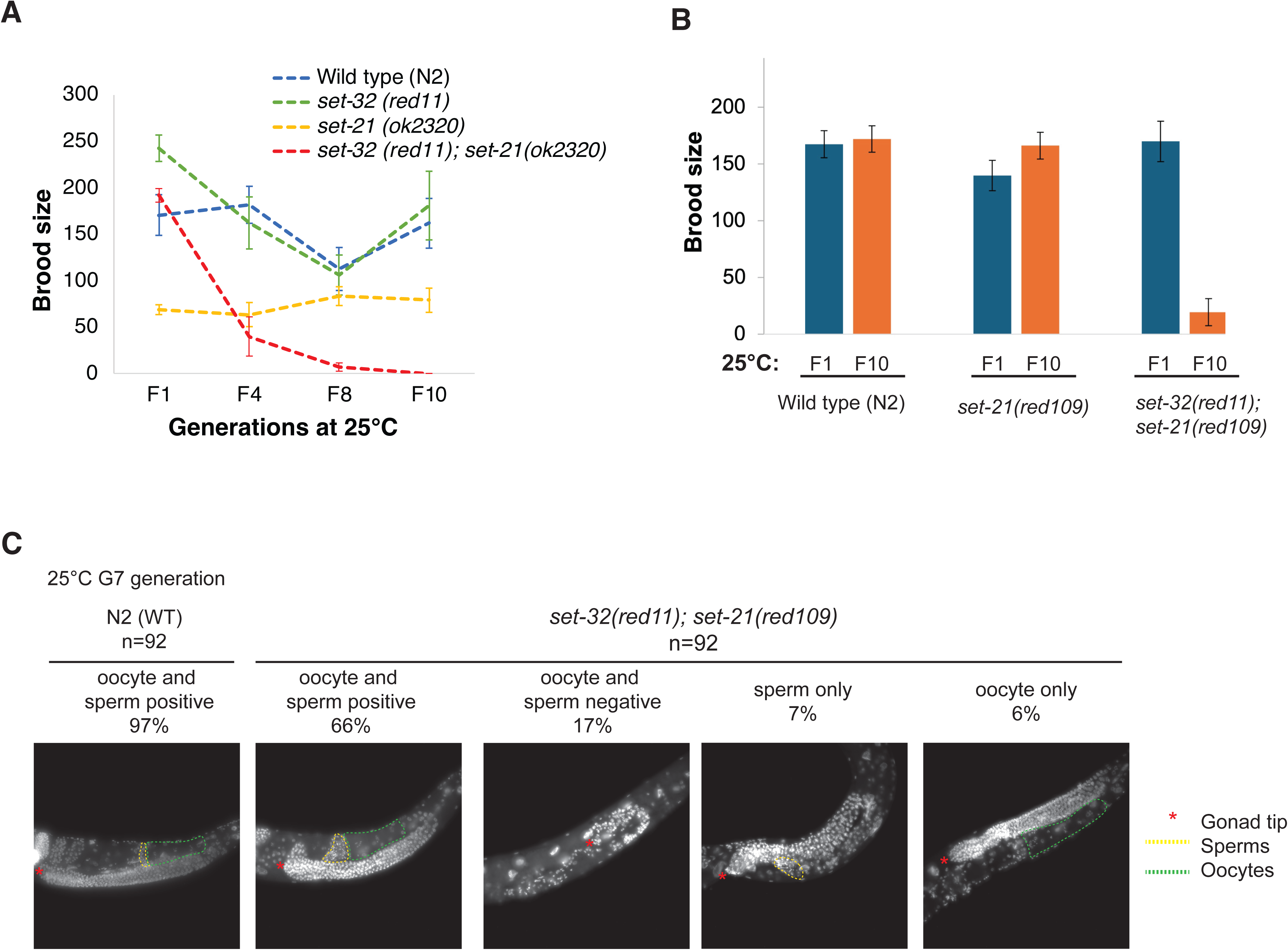
*set-32;set-21* mutant animals show germline defects at 25°C. (A-B) Multigenerational brood size analysis. Worms were maintained at 20°C before shifting to 25°C for F1 and the subsequent generations. Strains: WT (N2), *set-32(red11), set-21(ok2320)*, and *set-32(red11);set-21(ok2320)* mutant animals in (A) and WT (N2), *set-21(red109)*, and *set-32(red11);set-21(red109)* in (B). We note that the smaller brood size of *set-21(ok2320)* compared to *set-21(red119)* or *set-32;set-21(ok2320)* is likely due to some unknown background mutations. (C) Oocytes and sperm of *set-32(red11);set-21(red109)* young adults (F7 at 25°C) were examined by DAPI staining. Percentages of adult animals with both oocytes and sperm, only either oocyte or sperm, and neither gamete were indicated with representative DAPI-staining images.

**Figure S4.**
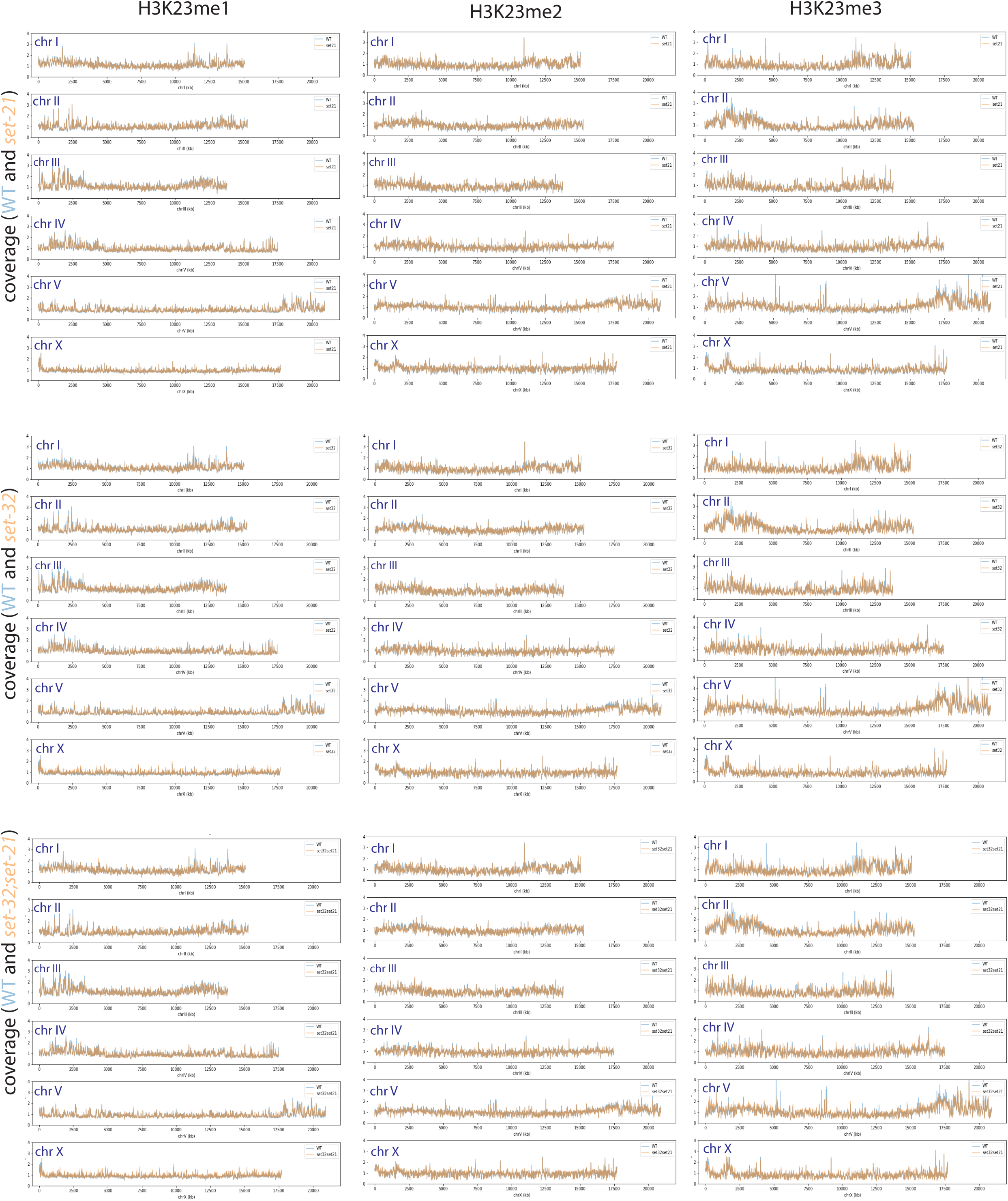
Whole-genome coverage plots of H3K23me1, me2, and me3 comparing WT versus *set-21*, *set-32*, or *set-32;set-21* mutant. The coverage, averaged from two replicates, was normalized to the ChIP input signal and was calculated for each 10kb window.

**Figure S5.**
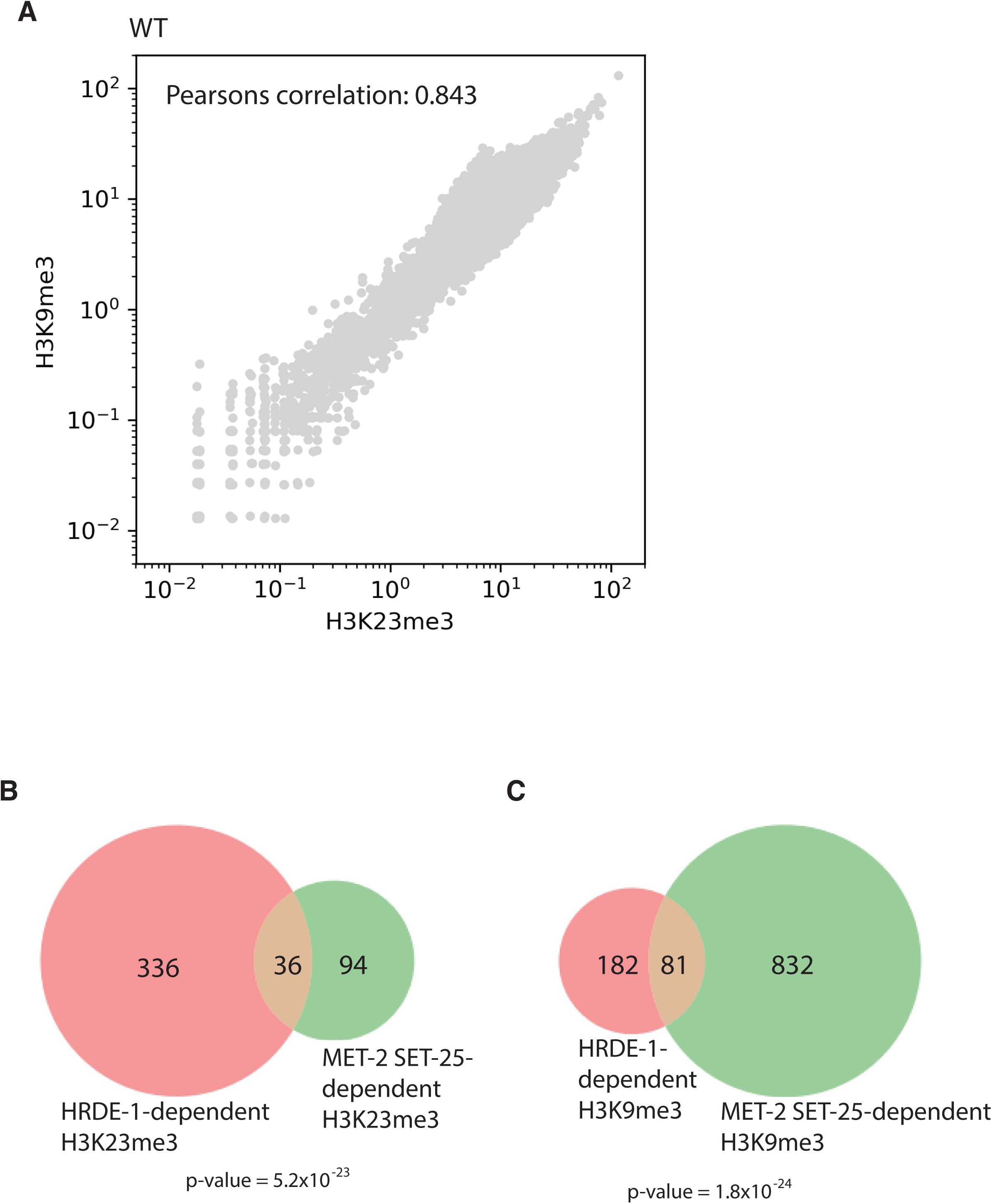
(A) A scatter plot of whole-genome comparison of H3K9me3 and H3K23me3 levels (1 kb windows) in the WT animals. (B) A Venn diagram of HRDE-1-dependent H3K23me3 and MET-2 SET-25-dependent H3K23me3. (C) A Venn diagram of HRDE-1-dependent H3K9me3 and MET-2 SET-25-dependent H3K9me3.

**Figure S6.**
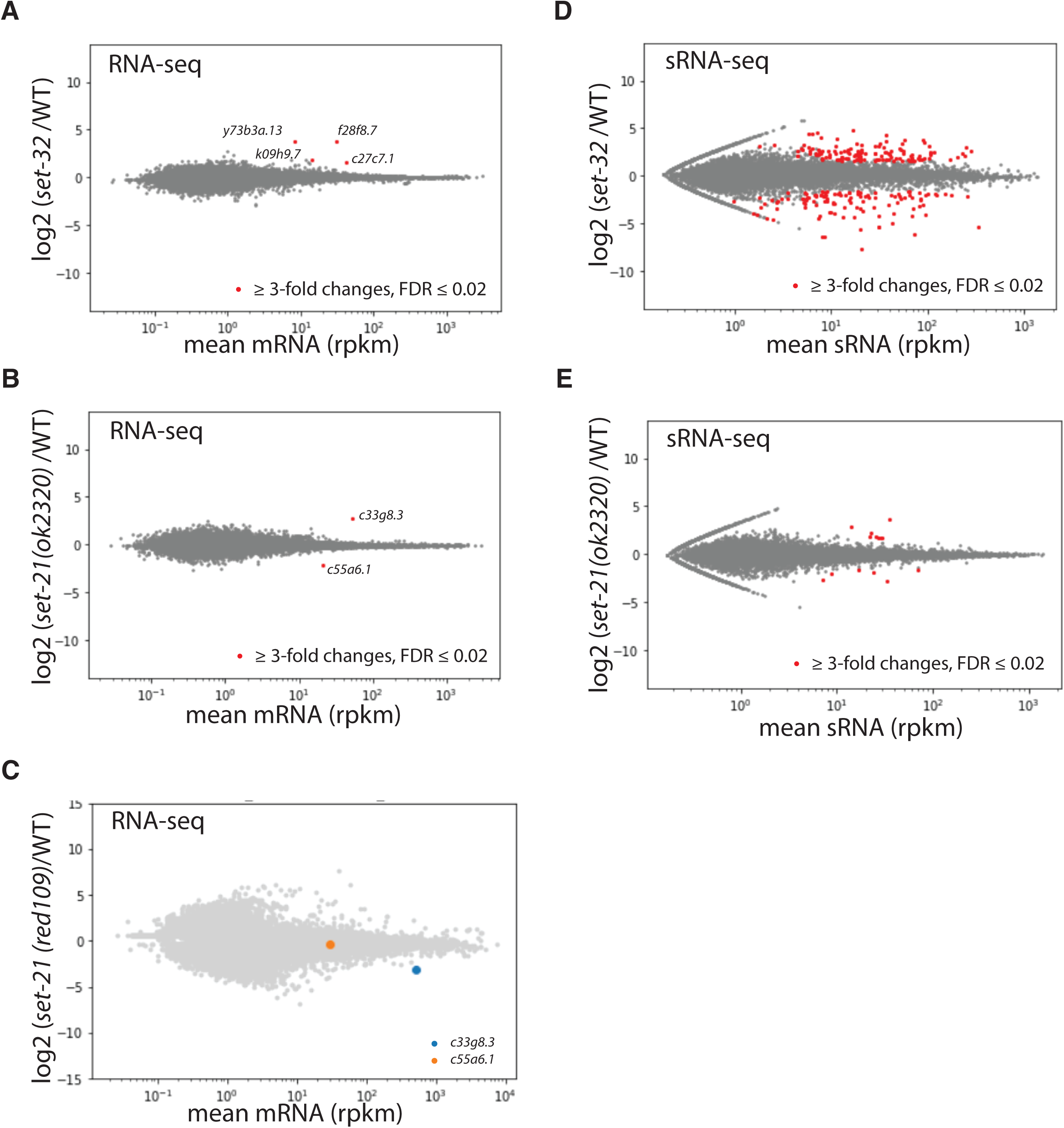
RNA-seq (A-C) and sRNA-seq (D-E) comparison of WT and *set-32* or *set-21* single mutant.

**Figure S7.**
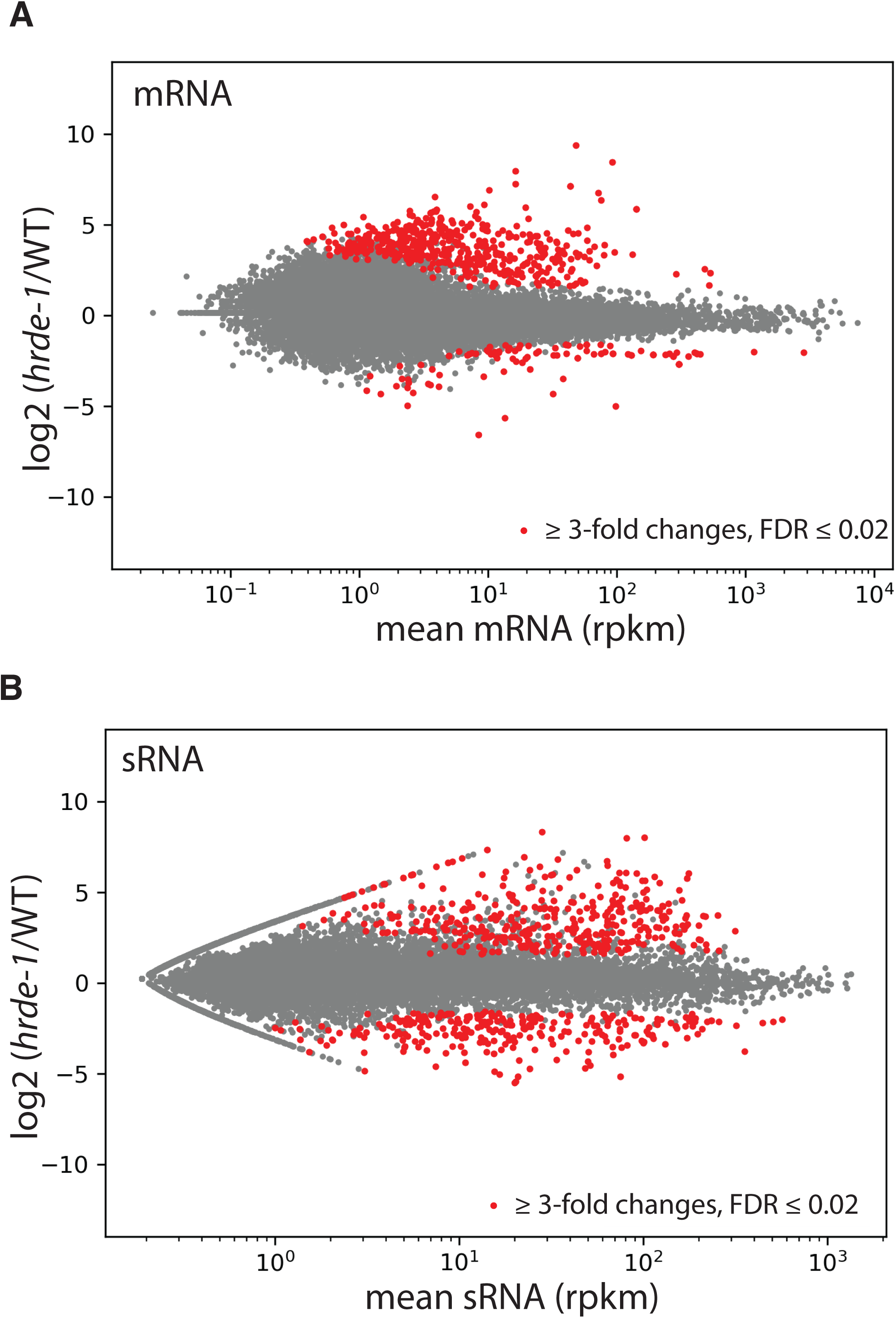
MA-plots comparing *hrde-1* and WT animals for (A) mRNA and (B) siRNA expressions of all protein-coding genes.

**Figure S8.**
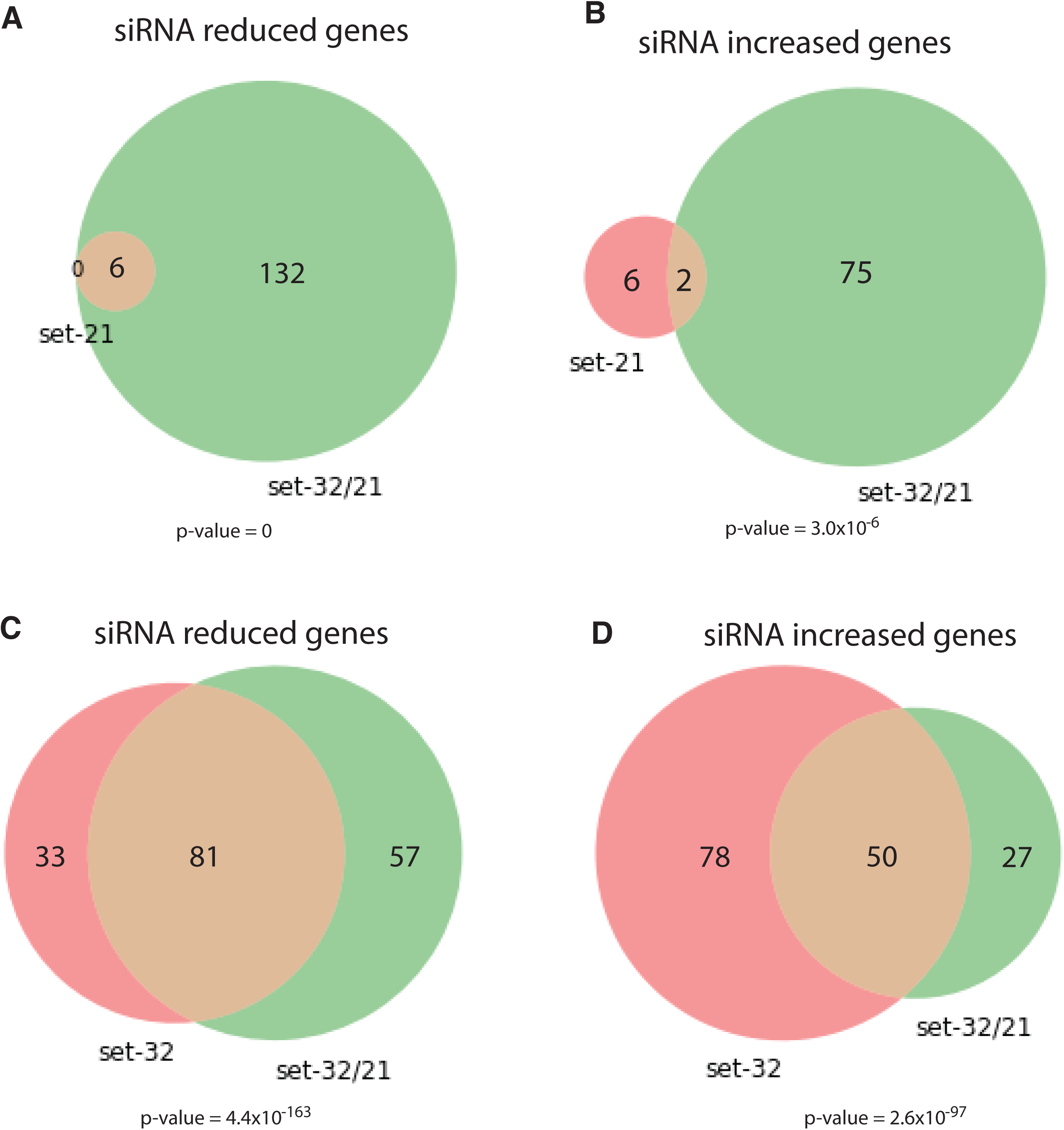
Venn diagram of genes with decreased or increased siRNA expression (minimal 3-fold change, FDR ≤ 0.02) comparing *set-21* or *set-32* single mutant with *set-32;set-21* double mutant.

**Figure S9.**
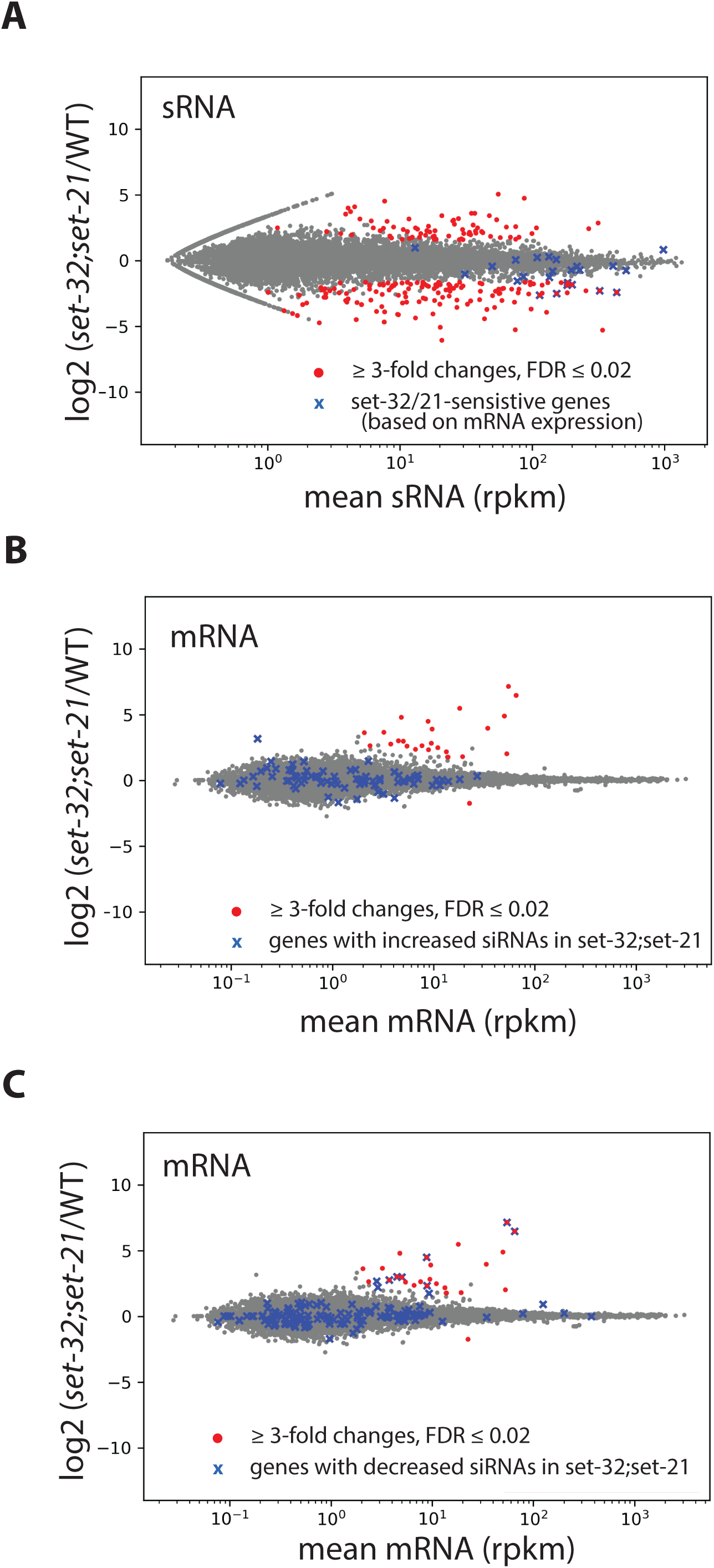
*set-32;set-21* mutations cause more wide spread changes in siRNA expression than changes in mRNA expressions. (A) sRNA MA-plot comparing *set-32;set-21* and WT with *set-32/21*-sensitive genes (based on mRNA-seq) highlighted. (B-C) mRNA MA-plots comparing *set-32;set-21* and WT with genes that had increased (B) and decreased (C) siRNA expression in the *set-32;set-21* compared to WT highlighted.

**Figure S10.**
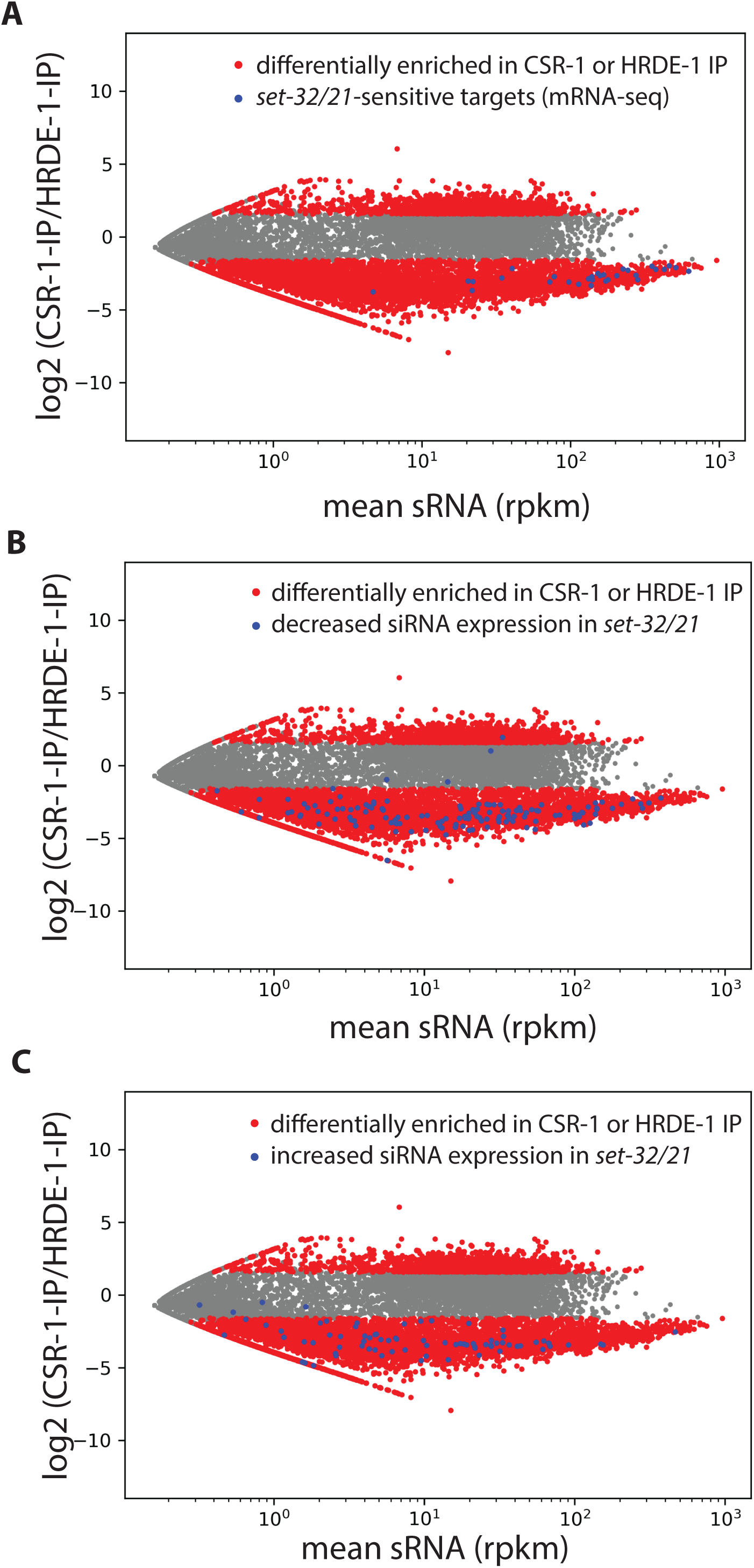
For genes of which mRNAs are desilenced in *set-32;set-21*, as well as genes of which siRNAs are differentially expressed in *set-32;set-21* (either decreased or increased), their siRNAs tend to be bound by HRDE-1, instead of CSR-1. The same CSR-1 vs HRDE-1-coIP sRNA MA-plot was shown in all three panels with each highlighting a different set of genes (marked in blue): (A) desilenced genes (mRNA-seq) in *set-32;set-21,* (B-C) genes with decreased (B) or increased (C) siRNA expression in the *set-32;set-21* mutant. CSR-1 vs HRDE-1-coIP sRNA data were from ^50^. Genes with a minimal of 3-fold difference in CSR-1-vs-HRDE-1-coIP siRNA (FDR≤0.02) were highlighted in red.

Table 1. A list of H3K23me3-enriched regions in WT identified in WT adult animals, with H3K23me3 ChIP-seq differential analysis outputs (log2 ratio, FDR and mean) for WT vs *hrde-1* and WT vs *set-32;set-21* comparisons calculated by BaySeq.

Table 2. Protein-coding gene differential analysis results of H3K23me3 ChIP-seq, Pol II ChIP-seq, RNA-seq, and sRNA-seq for the comparisons between WT and various mutant animals. Set-32/21-sensitive genes, based on RNA-seq analysis, were indicated.

Table 3. A list of high-throughput sequencing libraries used in this study.

